# Magnetoelectric materials for miniature, wireless neural stimulation at therapeutic frequencies

**DOI:** 10.1101/461855

**Authors:** Amanda Singer, Shayok Dutta, Eric Lewis, Ziying Chen, Joshua C. Chen, Nishant Verma, Benjamin Avants, Ariel K. Feldman, John O’Malley, Michael Beierlein, Caleb Kemere, Jacob T. Robinson

## Abstract

A fundamental challenge for bioelectronics is to deliver power to miniature devices inside the body. Wires are common failure points and limit device placement. On the other hand, wireless power by electromagnetic or ultrasound waves must overcome absorption by the body and impedance mismatches between air, bone, and tissue. In contrast, magnetic fields suffer little absorption by the body or differences in impedance at interfaces between air, bone, and tissue. These advantages have led to magnetically-powered stimulators based on induction or magnetothermal effects. However, fundamental limitations in these power transfer technologies have prevented miniature magnetically-powered stimulators from applications in many therapies and disease models because they do not operate in clinical “high-frequency” ranges above 50 Hz. Here we show that magnetoelectric materials – applied in bioelectronic devices – enable miniature magnetically-powered neural stimulators that can operate up to clinically-relevant high-frequencies. As an example, we show that ME neural stimulators can effectively treat the symptoms of a hemi-Parkinson’s disease model in freely behaving rodents. We further demonstrate that ME-powered devices can be miniaturized to mm-sized devices, fully implanted, and wirelessly powered in freely behaving rodents. These results suggest that ME materials are an excellent candidate for wireless power delivery that will enable miniature bioelectronics for both clinical and research applications.

## Introduction

Wireless neural stimulators have the potential to provide less invasive, longer lasting interfaces to brain regions and peripheral nerves compared to battery-powered devices or wired stimulators. Indeed, wires are a common failure point for bioelectronic devices. Percutaneous wires present a pathway for infection (Hargreaves et al. 2004) and implanted wires can also limit the ability of the stimulators to move with the tissue, leading to a foreign body response or loss of contact with the target tissue (Roy et al. 2007; Markwardt et al. 2013). Additionally, chronic stress and strain on wires, particularly for devices in the periphery, can lead to failure in the wire itself or its connection to the stimulator (Sahin & Pikov 2011). In small animals like rats and mice, wires used to power neural stimulators can interfere with natural behavior, particularly when studying social interaction between multiple animals (Pinnell et al. 2018).

One of the primary challenges for wireless neural stimulators is to create efficient miniature devices with cross sections roughly the size of a grain of rice (< 25 mm^2^ in area) that operate reliably beneath bone and tissue as an animal or human patient engages in normal activity. At areas of less than 25 mm^2^, devices could be fully implanted in the periphery and be light enough to allow for unrestricted animal behavior; however for devices this small, power delivery remains a challenge. Efficient power transfer with propagating electromagnetic waves requires antennas with feature sizes comparable to the electromagnetic wavelength. Thus, for sub-millimeter devices, such as the proposed RF powered “neurograins,”(Nurmikko 2018) effective power-transfer frequencies lie in the GHz range, where electromagnetic radiation is absorbed by the body (International & Safety 2006). Absorption of this radio-frequency electromagnetic energy limits the amount of power that can be safely delivered to implants deep inside tissue (International & Safety 2006). As a result, researchers typically turn to magnetic induction or batteries to power implanted devices; however, these techniques also limit the degree of miniaturization. Batteries increase the size of the device and add considerable weight. Additionally, batteries require replacement or charging, which can limit the potential uses. Inductively coupled coils, on the other hand, can be made smaller and lighter than batteries, however; the power a receiving coil can generate is directly related to the amount of flux captured by the area of the coils. Thus, when the receiver coils are miniaturized, the output power reduces and becomes more sensitive to perturbations in the distance or angle between the transmitter and receiver (Fotopoulou & Flynn 2011). For example, Freeman *et al.* demonstrated that small inductive coils less than 1 mm in diameter can power stimulators for the sciatic nerve in anesthetized rats (Freeman et al. 2017); however, the stimulation frequencies have been limited to less than 50 Hz and stable performance has been difficult to achieve in freely moving animals due to the reduced power coupling efficiency that accompanies changes in the angle and distance between the receiver and transmitter during behavior (Maeng et al. 2019). Near-field inductive coupling has also been used to power optogenetic stimulators in freely moving mice (Shin et al. 2017). While this approach is highly effective when the coils are at or near the surface of the skin, the relatively large cross-sectional area (70 mm^2^) of the receiving coil and high sensitivity to coil curvature could limit its application for bioelectronic implants that are deep inside the body.

Additionally, a number of therapeutic applications require high-frequency stimulation, which has been difficult to achieve with miniature bioelectronic devices. While some applications including vagus nerve stimulation and spinal cord stimulation can be effectively treated using low frequency (10-50 Hz) stimulation, many disorders like Parkinson’s Disease (PD), and obsessive-compulsive disorder, require stimulators in the high-frequency “therapeutic band” between 100 and 200 Hz (De Hemptinne et al. 2015; Alonso et al. 2015; Theodore & Fisher 2004). This type of high-frequency neural stimulation is challenging because charge on the electrode must be dissipated between successive stimulation pulses to prevent electrolysis, tissue damage, and changes to the local pH (Merrill et al. 2005). Charge dissipation at high frequencies is accomplished by using a biphasic stimulus waveform that actively or passively charges and discharges the electrode with each cycle. Indeed all clinically approved electrical neural stimulation therapies in this therapeutic band use various forms of “charge balanced” biphasic stimulation waveforms (Parastarfeizabadi & Kouzani 2017).

Recently, several new wireless power transfer technologies have enabled miniature neural stimulators; however, these approaches have yet to demonstrate high-frequency stimulation in freely moving animals. Montgomery et al. and Ho et al. have shown that one can use the mouse body as an electromagnetic resonant cavity to effectively transfer energy to sub-wavelength scale devices implanted inside the animal (Montgomery et al. 2015; Ho et al. 2015). This “mid-field” technique has also been further developed to use conformal phase surfaces to activate devices implanted in larger animals (Agrawal et al. 2017). This approach is particularly effective to drive tiny LEDs for optogenetic stimulation in mice. However, because the electrical waveform is monophasic, electrical stimulation has been limited to < 20 Hz and would require the use of active programmable integrated circuits to safely access higher frequencies. Using superparamagnetic nanoparticles to absorb energy from high-frequency (500 kHz) magnetic fields (Carrey et al. 2015), one can heat specific regions of the brain (Chen et al. 2015; Munshi et al. 2017) in freely moving animals (Munshi et al. 2017). This local heat can stimulate neural activity when the targeted brain region is genetically modified to respond to changes in temperature (Chen et al. 2015; Munshi et al. 2017). However this approach requires transgenesis, which adds regulatory complexity and has yet to show high-frequency operation due the requirement for the tissue to cool between stimulation pulses. Ultrasound provides a promising and efficient method to power bioelectronic implants because ultrasound wavelengths are 10^5^ times smaller than electromagnetic waves at the same frequency allowing sub-millimeter-sized devices to have wavelength-scale piezoelectric antennas (Johnson et al. 2018; Seo et al. 2016; Piech et al. 2020). However, implementation of these “StimDust” motes can be challenging in freely moving animals because the impedance mismatch between air, bone, and tissue requires contact between soft tissue and the ultrasound transducer for efficient power transfer. As a result, there has yet to be a demonstration of ultrasound-powered neural stimulators in freely moving animals. For a more detailed comparison of these current technologies see Table 1.

**Table 1.** Comparison of miniature neural stimulators.

| Technology | Inductive Optogenetics | E-Particle | Mid-Field | Mid-Field | Stim Dust | RF Antenna | Magneto thermal 1 | Magneto thermal 2 | This work demo 1 | This work demo 2 |
| --- | --- | --- | --- | --- | --- | --- | --- | --- | --- | --- |
| Reference | [a] | [b] | [c] | [d] | [e] | [f] | [g] | [h] | This work | This work |
| Power Type (Implant) | Inductive Coil | Inductive Coil | Mid-Field Coil | Mid-Field Coil | Piezo electric | RF antenna | Nano-particle injection | Nano-particle injection | Magneto electric | Magneto electric |
| Activation by | Magnetic Field | Magnetic Field | Electro magnetic | Electro magnetic | Ultra-sound | Electro magnetic | Magnetic Field | Magnetic Field | Magnetic Field | Magnetic Field |
| Carrier Frequency | 13.56 MHz | 10.9 MHz | 1.5 GHz | 1.6 GHz | 1.85 MHz | 2.3-3.2 GHz | 500 kHz | 500 kHz | 100-170 kHz | 250-400 kHz |
| Stimulation Type | LED | Electical | LED | Electrical | Electical | LED | Heat | Heat | Electrical | Electrical |
| Whole Implant Size (mm<sup>3</sup>) | 100 | 0.5 | 10-25 | 15 | 6.5 | 24 | N/A | N/A | 500 | 175 |
| Size of Power Source (mm<sup>3</sup>) | 40 | 0.25 | 5 | 5 | ~0.5 | 20 | N/A | N/A | 20 | 2-4 |
| Implant Weight (mg) | 30 | - | 25-50 | 70 | 10 | 33 |  |  | 500 | 500 |
| Max Implant Power (in animal, mW) | 30 | 0.036<sup>j</sup> | 10 | 2 | 0.35 | - | N/A | N/A | 0.1-0.2 | 2 |
| Demonstrated Charge Balance (y/n) | N/A | no | N/A | no | yes | N/A | N/A | N/A | yes | yes |
| Freely Moving (y/n) | yes | yes | yes | no | no | yes | no | yes | yes | yes |
| Arena Volume (cm<sup>3</sup>) | 12000 | 1000<sup>j</sup> | 1000 | N/A | N/A | video tracked area | N/A | 300 | 3000-7000 | 500 |
| Magnetic Field Strength (mT) | - | 0.04 | - | - | - | - | 19 | 9.4 | 1-2 | 1-2 |
| Genetic Modification (y/n) | yes | no | yes | no | no | yes | yes | yes | no | no |
| Fully Implanted (y/n) | yes | yes | yes | yes | yes | yes | yes | yes | no | yes |
| Stimulation Frequency (Hz) | - | 50 | - | ~3 | 2 kHz | - | < 1 (0.02) | < 1 (0.005) | 200 | 150 |
| Average Power Required (W) | - | - | 4 | - | - | - | - | 7500 | 30 | 7 |

<sup>j</sup>estimated based on published data
References: [a](Shin et al. 2017)[b](Freeman et al. 2017; Maeng et al. 2019) [c] (Montgomery et al. 2015) [d] (Ho et al. 2015)[e](Piech et al. 2020) [f] (Park et al. 2016) [g] (Chen et al. 2015)[h] (Munshi et al. 2017)

Here we show that magnetoelectric (ME) materials enable magnetically powered miniature neural stimulators that operate across the low- and high-therapeutic frequencies. Similar to inductive coils, these materials transform a magnetic field to an electric field, but instead of using an implanted coil we use a material that generates a voltage via mechanical coupling between magnetostrictive and piezoelectric layers in a thin film. Namely, the magnetic field generates strain in the magnetostrictive layer as the magnetic dipoles align with the applied field. That strain exerts a force on the piezoelectric layer, which generates a voltage (Fig. 1). By exploiting this transduction mechanism, magnetoelectrics do not suffer from the same miniaturization constraints as coils and can be driven by weak magnetic fields on the order of a few millitesla. These properties have led researchers to propose magnetoelectrics as a promising material for bioelectronic implants (Nan et al. 2017; O’Handley et al. 2008; Yue et al. 2012; Guduru et al. 2015; Ribeiro et al. 2016). Here we demonstrate proof-of-principle wireless neural stimulators based on ME materials and show that they effectively treat the symptoms of Parkinson’s Disease (PD) in a freely behaving rodent disease model. We further show that ME materials can be miniaturized to the size of a grain of rice, be fully implanted under the skin, and effectively stimulate the brain of freely behaving rodents to drive a place preference.

**Figure 1.**
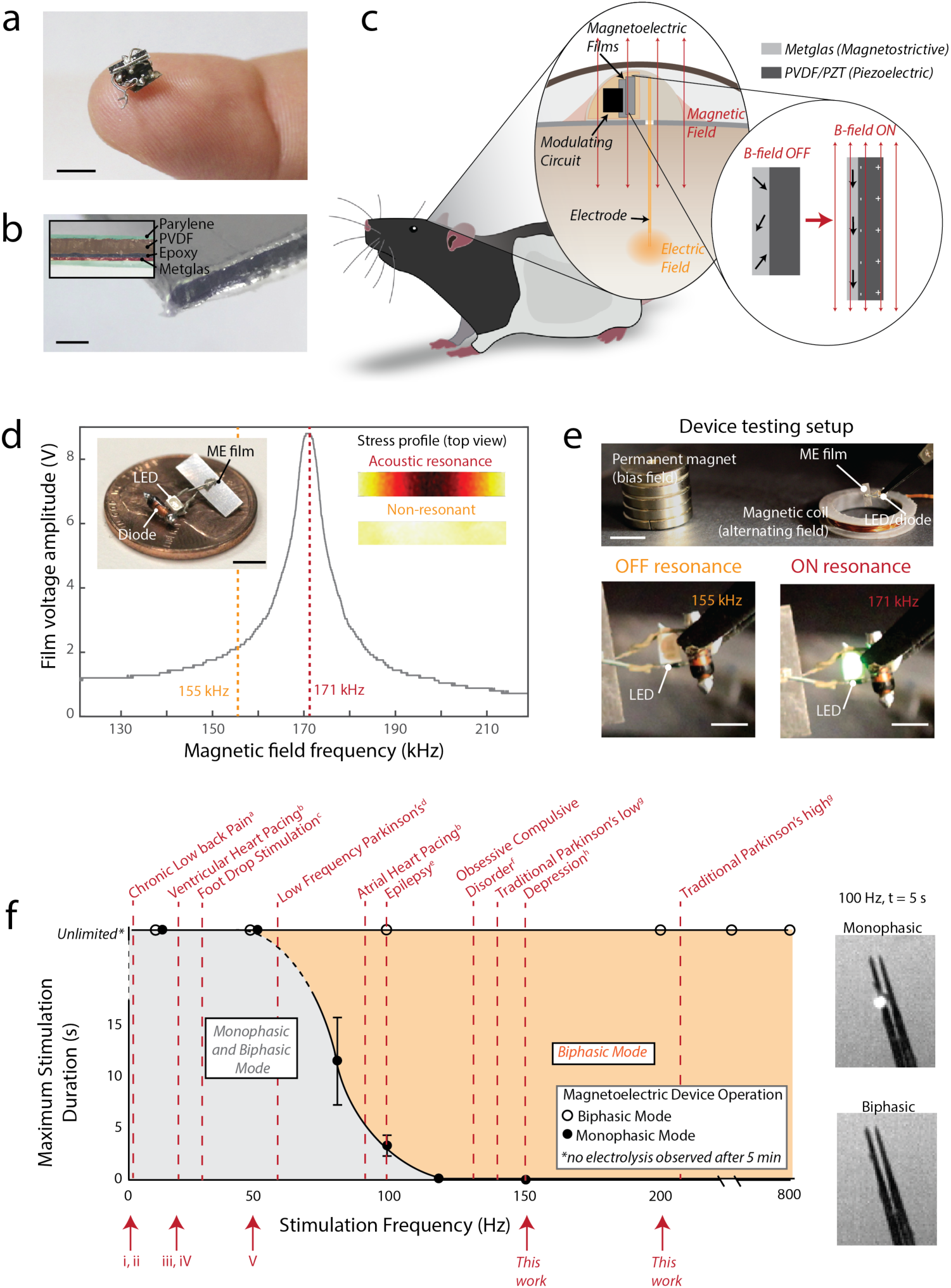
ME films convert alternating magnetic fields into a voltage. (a) Photo of a mm scale ME device shown on a fingertip scale bar = 5 mm (b) Cross sectional image of a cut ME film showing the layers of the laminate, scale bar = 0.2 mm (c) Diagram of a ME device on a freely moving rat for wireless neural stimulation. The active ME element consists of piezoelectric PVDF or PZT (dark grey) and Metglas (light) laminate encapsulated by Parylene-C. Inset shows the operating principle whereby the strain produced when magnetizing the light gray magnetostrictive layer is transferred to the dark grey piezoelectric layer, which creates a voltage across the film. (d) Example of a resonant response curve for a ME film showing that the maximum voltage is produced when the magnetic field frequency matches an acoustic resonance at 171 kHz. Photograph inset shows an example of an assembled ME stimulator. The “Stress profile” inset shows a top view of the stress produced in a ME film as calculated by a finite element simulation on and off resonance (COMSOL). (e) Device testing setup with a permanent magnet to apply a bias field and an electromagnetic coil to apply an alternating magnetic field (scale bars: upper = 1 cm, lower = 2 mm) (f) Maximum stimulation duration (using a 400 μs/phase pulse repeated at increasing frequencies) for a ME device in biphasic and monophasic operation. Maximum stimulation time is determined by time of electrolysis on a stereotrode in saline as evidenced by gas bubbles (error bars +/- 1 standard deviation for n=4 trials). Dashed red lines indicate frequencies of electrical stimulation used in various clinical applications, showing that biphasic operation is necessary for many clinically relevant applications (a: (Schabrun et al. 2014) b: (Mulpuru et al. 2017) c: (Kesar et al. 2010) d: (Baizabal-Carvallo & Alonso-Juarez 2016) e: (Theodore & Fisher 2004) f: (Alonso et al. 2015) g: (De Hemptinne et al. 2015) h:(Bewernick et al. 2010)). Roman numerals indicate stimulation frequencies demonstrated by previously published miniature magnetic stimulators (i: Magnetothermal, Chen et. al, 2015, ii: Magnetothermal, Munshi et. al, 2017, iii: Mid-Field Optogenetics, Montgomery et. al, 2015, iv: RF Inductive Coupling, Freeman et. al, 2017, v: RF Inductive Coupling, Maeng et. al, 2019).

## Results

### Fabrication and characterization of ME stimulators

We fabricated proof-of-principle miniature ME stimulators (Fig. 1a,c) by bonding a rectangular magnetostrictive layer (Metglas) to a platinum coated piezoelectric layer, polyvinlydine fluoride (PVDF) or lead zirconate titanate (PZT). We then encapsulated the films in a protective parylene-C layer (8-10 μm thick) (Fig. 1b, see Methods). We used PVDF or PZT layers between 28 and 110 μm, which yielded total device thicknesses between 50-150 μm. When we measured the voltage across the film, we found a dramatic voltage increase when the applied magnetic field frequency, matches an acoustic resonant frequency (Fig. 1d). Because the resonant frequency is proportional to the inverse of the film length (Fig. S1i), we can design multiple films and selectively activate them by changing the stimulus frequency. Using this principle, we can use different magnetic field frequencies to activate separate monophasic devices that may be in different areas of the body, or create biphasic stimulators by interleaved resonant stimulation of two different films, with each film driving either the positive or negative phase of the neural stimulus.

We can further enhance the voltage generated by the ME films by applying a constant bias field with a permanent magnet or an electromagnet (Fig. 1e). Because the strain in the magnetostrictive material is a sigmoidal function of the magnetic field strength, the change in voltage produced by the alternating field is largest when the field oscillates about the midpoint of the sigmoid (Fig. S1a-c)(Zhai et al. 2006; Kulkarni et al. 2014). Thus, we use a bias field to offset the magnetic field near the center of the sigmoidal magnetostrictive response curve (approximately 8-9 mT for devices used here). This bias field allows us to generate therapeutic voltage levels while applying a small (few mT) alternating magnetic field around this central bias point using an electromagnetic coil and custom control circuitry that specifies the frequency and timing of the alternating magnetic field (Fig. S1e-h). Our references to magnetic field strength in this work refer to the amplitude of the alternating magnetic field around this bias point.

To identify the safe operational conditions for our ME stimulators we tested them in saline over a range of stimulation frequencies. We found that a biphasic stimulation waveform allowed us to apply constant stimulation up to at least 800 Hz without significant hydrolysis and monophasic stimulation could be safely applied up to approximately 50 Hz. For these tests we used stimulation amplitude of 2 V and a duration of 400 μs/phase (which is common for in vivo experiments). These ME films were attached to a stereotrode (Microprobes) in saline (see Methods) and the safe ranges were determined by measuring the time at which we could see bubbles at the electrode tip (Fig 1f) resulting from hydrolysis. This hydrolysis event indicates conditions that would lesion the surrounding tissue. We found that with a monophasic stimulation waveform, stimulation frequencies above 50 Hz produced hydrolysis while biphasic charge-balanced stimulation showed no hydrolysis up to the maximum tested frequency of 800 Hz. Compared to previously demonstrated miniature magnetic neural stimulators that operated in a monophasic stimulation mode, the biphasic ME devices shown here can access the high-frequency bands used for clinical applications like the treatment of Parkinson’s disease and obsessive-compulsive disorder (Fig 1f). The ME devices can also operate safely in a monophasic mode for low-frequency applications such as heart pacing or chronic pain stimulators, which can be achieved with a simplified device with a single ME film. One should note that the exact safety windows will depend on the amplitude, duty cycle, and electrode configuration of the stimulator.

An additional challenge for any wirelessly powered neural stimulator is to maintain a well-regulated stimulation voltage. This challenge is especially prevalent as devices become small, which often reduces the power transfer efficiency resulting in a greater sensitivity to the alignment between the device and power transmitter. ME materials offer two main advantages that can enable stable and effective stimulation even as devices become small and move with respect to the driver coils:

First, ME devices generate voltages well in excess of the effective stimulation potential, allowing them to be effective even if the materials are misaligned with the driver coils. At resonance, we have measured open-circuit ME voltages in excess of 30 V at a field strength of only 1 mT (Fig. S1k, m). Because effective stimulation voltages are usually between 1-5 V, we can cap the applied voltage to this effective stimulation range using an LED or Zener diode. As long as the voltage generated by the ME film is greater than or equal to the capping voltage, our device could apply the same stimulus voltage regardless of the angle or distance between the driver coil and the ME film. For a typical film we found that we could reorient the film by +/- 80 degrees and maintain voltages in excess of 3 V (Fig. S1d). This large angular tolerance is aided by the large magnetic permeability of the Metglas layer, which helps to direct the magnetic field lines along the long axis of the film, where they are most effective at creating a magnetostrictive response.

Second, the voltage generated by a piezoelectric material depends on the thickness of the piezoelectric layer and not the area of the film(Wan & Bowen 2017), allowing us to fabricate small magnetoelectric films that generate roughly the same stimulation voltage as larger devices. Figure S1j-k shows the peak voltage generated and quality factor for ME films of different areas. We found that, for a 52 μm thick PVDF layer, the voltage remains around 10 V even as the film length decreases. Variations of +/- 40 % in peak voltage and quality factors are likely due to defects produced during film fabrication, which may be reduced with improved manufacturing. We also verified that the output voltage depends only on the piezoelectric film thickness by measuring the peak voltages from ME devices with three different thicknesses of PVDF: 28 μm, 52 μm, and 110 μm. As expected, we see that the peak voltage increases linearly with the PVDF thickness and is independent of the film length.

For applications such as current delivery through implanted electrodes, where the available power is an important figure of merit, the advantage of our ME technology is the power we get from a mm-sized magnetically powered device. This allows us to perform experiments that require high power like high-frequency biphasic stimulation. Our calculations and experimental data show that the power generated by a ME device is proportional to the film area for a given thickness and a length-to-width ratio >3 (see Fig. S1l). This output power is also dependent on the *d*_*31*_ coefficient of the piezoelectric material (Fig. S1). This coefficient is 22 pC/N for PVDF (Precision Acoustics) and 320 pC/N for PZT (Piezo Systems). Despite the decrease in power as films become smaller, we calculate that PVDF/Metgals films less than 10 mm long can generate up to 4 mW, which is sufficient for many wireless applications including neural stimulation (Amar et al. 2015). In applications requiring higher power at miniature scales, we use PZT/Metglas ME films due to the higher *d*_*31*_ of PZT.

### Monophasic stimulation by ME films modulates cellular activity in vitro

Using fluorescence microscopy to image voltage in cultured cells, we found that monophasic stimulation for 50 ms at 100 Hz directly from PVDF/Metglas ME films reliably stimulated action potentials (APs). For these experiments we used “spiking” human embryonic kidney (HEK) cell lines that were modified to express sodium and potassium channels (see Methods). These cells have spike-like electrical waveforms that are rectangular in shape and can last for a few seconds depending on the confluency of the culture (Park et al. 2013). To determine the relative timing between magnetic stimulation and action potential generation, we transfected these cells with ArcLight (Jin et al. 2012)-a genetically encoded voltage indicator that allows us to measure action potentials using fluorescence microscopy.

To image fluorescence while we applied magnetic fields, we developed an experimental setup that allows us to place cells and ME films beneath an objective lens at the center of a 10 cm long solenoid with a 3 cm gap in the center. This configuration allowed us to place ME films, cells, and the objective lens at the center of the applied magnetic field (Fig. 2a). The field strengths (<1 mT, Fig. S2f) and frequencies (20-40 kHz) used in this experiment did not produce noticeable heat in our metallic objective lens or interfere with our imaging. Two slightly larger coils placed on either side of the gap provided the constant bias field.

**Figure 2.**
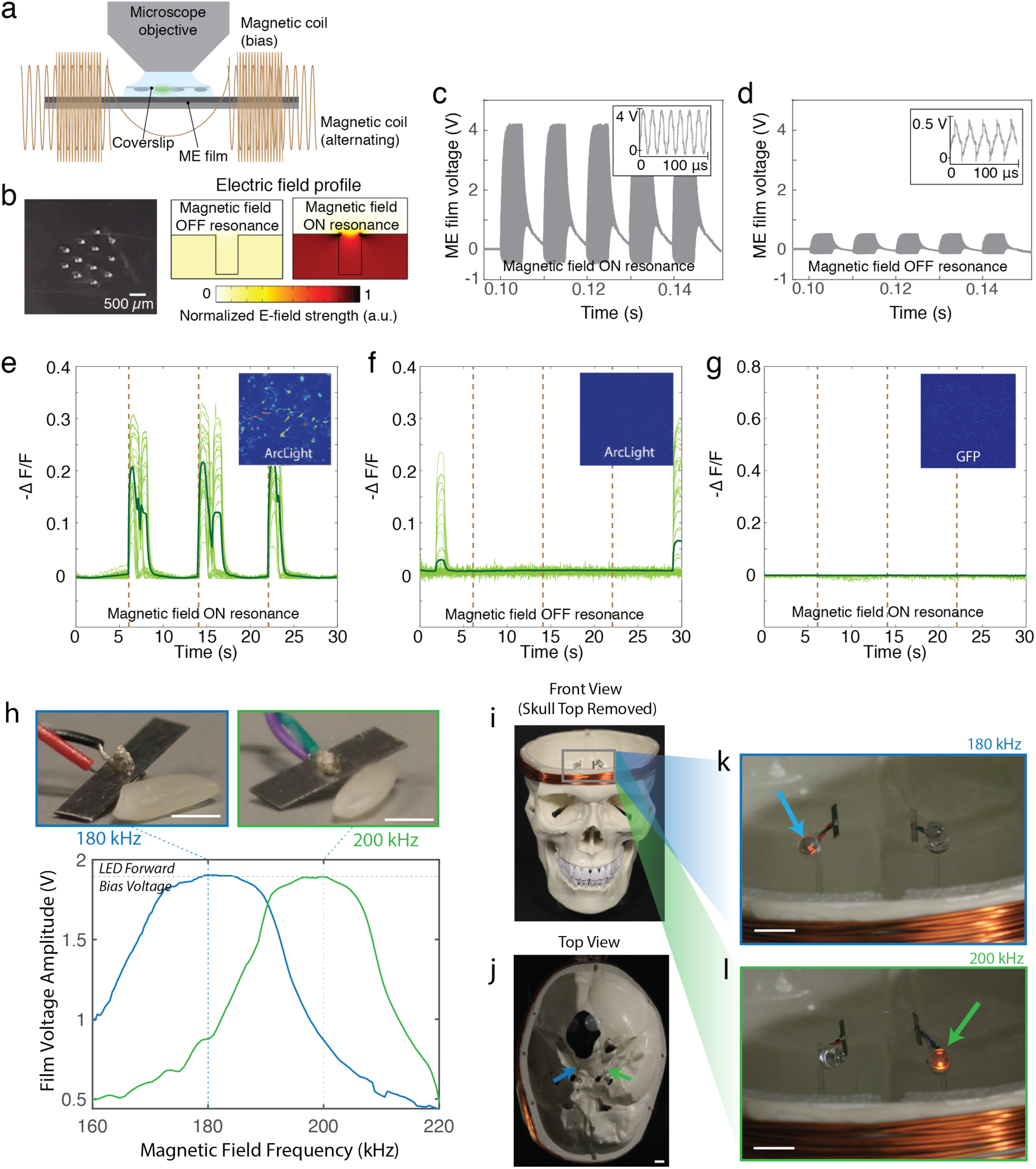
Monophasic ME stimulators activate cells in vitro. (a) Schematic of the experimental setup (b) Microscope image of holes stamped into the ME film and finite element simulation of the electric field shows that the holes produce fringing electric fields that overlap the culture cells (c) Voltage across the ME film when the magnetic field is on resonance and (d) off resonance. Insets show a zoom in of the high frequency carrier waveform. (e-g) Fluorescence from spiking HEKs transfected with ArcLight show action potentials are triggered by the ME film driven at resonance (e), but not when the film is driven off resonance (f). Fluorescence from HEK cells transfected with GFP (g) show no response when the ME film is driven on resonance confirming that the measured ArcLight response is the result of a change in transmembrane potential and not an artifact of the magnetic field or acoustic resonance of the ME film. (h) Photos of miniature ME films next to a grain of rice and the corresponding voltage as a function of magnetic field frequency (field strength 0.5 mT, scale bars 2 mm) (i) Front view and (j) top view of skull phantom with the top removed to view LEDs (film locations indicated by arrows, scale bar 1 cm) (k) Photo of LEDs attached to ME films with the magnetic fields at applied at 180 kHz and (l) 200 kHz. Selective illumination of the LEDs corresponding the resonant frequencies of the films demonstrates successful multichannel activation of individual films (scale bars 1 cm). Magnetic field strength was measured to be 0.5 mT at the location of the ME films.

We then approximated an implanted ME stimulator using two experimental configurations: 1) growing cells directly on the ME film (Fig. S2) and 2) laying a coverslip with adherent cells on top of the ME film (Fig. 2). To culture cells directly on the ME film, we coated the top parylene layer with poly-l-lysine. The healthy proliferation of HEKs on the ME device indicates that this encapsulation approach prevents the ME materials from limiting cell growth (Fig. S2b). However, in a typical use case, the target cells may not adhere to the ME stimulator, so we also tested the response of cells laid on top of the ME materials. In this configuration we first grew the cells on coverslips for 3-5 days before inverting the coverslips and laying them on the ME for testing (Fig. 2, see Methods).

To create fringing electric fields that interact with the cultured cells, we stamped holes in the ME film (Fig. 2b). The films were otherwise fabricated as described above (Fig. 1, Methods). In experiments using ME films and Pt electrodes we found that high-frequency biphasic stimulation at the ME resonance frequency (typically 20-150 kHz) was not effective to stimulate APs in cultured HEKs, as predicted by the low-pass filtering properties of the cell membrane (Freeman et al. 2017). To create an effective monophasic stimulus waveform, we used a Schottky diode to rectify the voltage to create entirely positive or negative voltage waveforms depending on the diode direction. This rectified waveform has a slowly varying monophasic envelope in the <500 Hz frequency band where cells are responsive (Fig. 2c,d).

For both cells grown directly on the ME films and those placed in contact we found that five stimulation pulses with an envelope frequency of 100 Hz (applied for only 50 ms total so as not to introduce hydrolysis) consistently stimulated APs in the spiking HEK cells (Fig. 2e, S2d, Supplementary Video 1). Critically, this stimulation frequency is within the therapeutic window for many deep brain stimulation treatments (So et al. 2017), and difficult to achieve with other wireless stimulators that compensate for low-efficacy energy harvesting by charging on-board capacitors (Sun et al. 2017). For our experiments, the carrier frequency of the magnetic field was at the resonant frequency of the device, which varied between 20-40 kHz depending on device length (Fig S2c). To test stimulation reliability, we repeated the 5-pulse stimulus three times over a period of 30 seconds. We observed APs for each stimulation pulse in n = 43 cells on coverslips and n = 144 cells grown on films. In these experiments all cells in the field of view were activated simultaneously due to the fact that HEK cells are known to be electrically coupled when grown to confluence. We confirmed that the APs stimulated by the ME film were in fact the result of resonant excitation of the film and not an artifact of the applied magnetic fields by imaging voltage-sensitive fluorescence when the magnetic field was tuned off of the resonant frequency. For non-resonant excitation we observed no correlation between the applied field and fluorescently detected APs in the spiking HEKs (Fig. 2f, S2e), supporting the conclusion that APs were stimulated by the ME film at resonance. We also confirmed that the fluorescent signal recorded indeed represents the voltage-dependent ArcLight response by imaging cells transfected with voltage-independent cytoplasmic GFP. These cells showed no change in fluorescence when the films were driven at the resonant frequency (Fig. 2g). This type of monophasic stimulation has applications such as low frequency deep brain stimulation for movement disorders (Baizabal-Carvallo & Alonso-Juarez 2016).

### ME devices can be individually addressed based their resonant frequency

Another advantage of ME technology is that we can individually address multiple implanted devices by fabricating the films to have unique resonant frequencies. Furthermore, we can miniaturize the device while maintaining effective stimulation capabilities because the voltage of the ME stimulators depends on the thickness of the piezoelectric layer (not the area of the film). As a proof-of-concept demonstration we show that two rice-sized ME films with cross sectional areas of ∼16 mm^2^ can be individually addressed at the center of a human skull phantom using an external electromagnet. These two-films with lengths of 8 mm and 10 mm have acoustic resonant frequencies of 180 and 200 kHz (Fig. 2h). When these films are attached to an orange LED, their output voltage is capped at approximately 1.8 V, which helps to regulate the stimulation voltage and allows us to visualize film activation. This choice of capping voltage could be modified to match the target stimulation voltage. ME films of this size are smaller than current DBS leads and could potentially be implanted into deep brain areas as shown in Fig 2i-l. Additionally, the magnetic stimulation coil is small enough to be incorporated into a stylish hat or visor that could be worn comfortably by a patient. When we placed the two ME films at the center of a skull phantom we found that we could individually illuminate the LEDs on each film when we applied a magnetic field at the resonant frequency of the selected film (Fig 2i-l). For this experiment we used a 40 W power supply, which produced a field of approximately 0.5 mT at the center of the skull phantom. The top of the skull phantom was removed for visualization, but had no effect on our ability to drive the LED indicators. The number of stimulation channels could be increased with the addition of ME films with different resonant frequencies.

This technology enables independent external wireless control of multiple miniature stimulators deep beneath a human skull phantom. As mentioned above, radio-frequency (RF) powered antennas that operate at frequencies above ∼1 MHz have limitations in the amount of power that can safely be delivered to an implanted device without causing potentially harmful tissue heating. Simulations show that when operating with the safe power limits, RF-antennas must be placed on the surface of the brain or in very shallow regions to harvest sufficient power for neural stimulation. “Mid-field” techniques (Agrawal et al. 2017), improve the RF coupling efficiency enabling deep operation, but because this approach operates at a fixed frequency, individually addressable motes have yet to be demonstrated. Other techniques for wireless power delivery discussed previously, such magnetic induction, also cannot achieve deep, miniature, multichannel stimulation.

### Biphasic stimulation from ME films can drive high-frequency neural activity in vitro

As described above, biphasic stimulation is used for most biomedical stimulators because it can create a charge-balanced stimulus that reduces charge buildup and undesired electrochemical reactions at the electrode surface(Merrill et al. 2005). While the voltage waveform produced by ME films is biphasic at the resonant frequency (typically 20 – 150 kHz), these frequencies are too high to produce reliable cell stimulation. To create an effective biphasic stimulus in the therapeutic window (100 – 200 Hz), we use two films with distinct resonant frequencies connected to the same stimulating electrodes (Fig. 3a,c). The first film is attached to a full wave rectifier, which is oriented to generate a positive pulse, while the second film is attached to a full wave rectifier that generates a negative pulse. These transistors block currents generated by one film from propagating through the circuitry attached to the other film, ensuring that only one half of the circuit is active at a time. By switching the magnetic field frequency between the two ME resonant frequencies, we can alternate positive and negative phase stimulation to create a biphasic neural stimulator (Fig. 3d,e). In this case the residual charge of <1 nC, which discharges in <2 ms, implies that this stimulator can safely operate at frequencies up to >500 Hz without accumulating charge.

**Figure 3.**
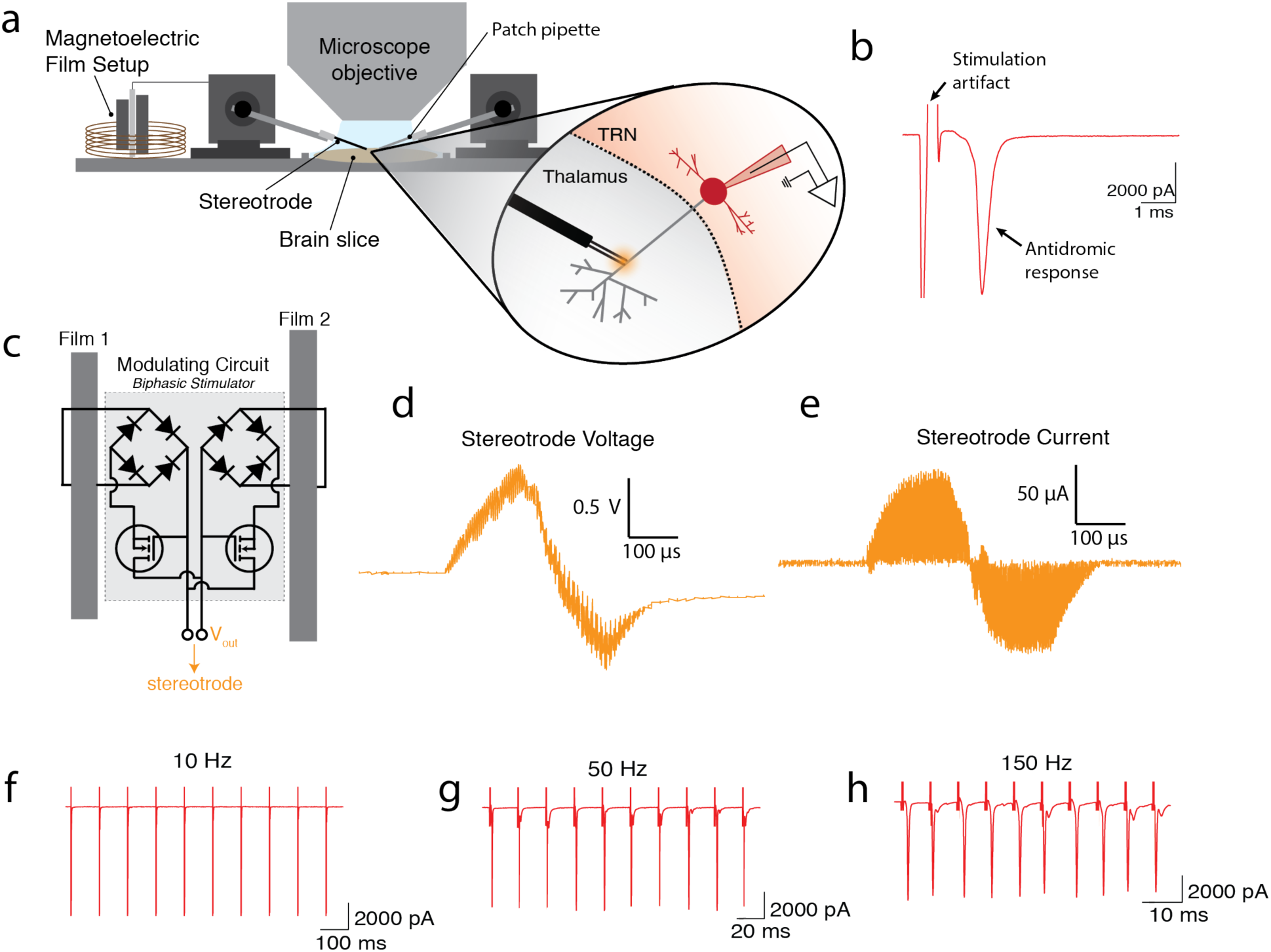
Biphasic ME stimulators activate neurons in ex vivo brain slices. (a) Schematic of experimental setup with two ME films for biphasic stimulation. A bipolar stereotrode was placed into the ventrobasal nucleus of the thalamus to activate axons of TRN neurons (b) Representative voltage-clamp recording in TRN showing stimulation-triggered short-latency antidromic spike (artifact cropped for scale). (c) Schematic of the circuit used to generate the biphasic waveform (d) Measured voltage across the stereotrode shows the expected biphasic pulse shape (e) Calculated current based on measuring the voltage across a load resistor shows nearly perfect charge balancing with <1 nC accumulating on the electrode per pulse train. (f-h) Recorded spike activity at various frequencies of ME stimulation demonstrates the ability of the ME device to reliably entrain action potential activity at 10, 50, and 150 Hz.

We demonstrated that clinically relevant regions of high frequency stimulation are safely accessible with this ME powered device by using our biphasic ME stimulator to drive high-frequency neural spikes in a mouse brain slice (see Methods). We stimulated axons of thalamic reticular nucleus (TRN) neurons by placing a stereotrode attached to the ME stimulator into the adjacent ventrobasal nucleus of the thalamus and performing whole-cell recordings in TRN neurons. (Fig 3a). We found that short-latency antidromic spikes were reliably evoked (Fig 3b), with the recorded spike frequency matching the programmed magnetic field envelope frequency (10, 50, and 150 Hz) confirming that neuronal activity can be precisely controlled using our ME stimulation (Fig 3f-h).

We also found that our biphasic ME stimulator is capable of repeatable neural stimulation using neocortical brain slices derived from mice that express the genetically encoded calcium indicator GCaMP3 in glutamic acid decarboxylase 2 (GAD2) positive GABAergic neurons (Fig. S3). To image neural activity following ME stimulation we again inserted a stereotrode attached to the biphasic ME stimulator described above while we imaged GCaMP activity using fluorescence microscopy (Fig. S3a-c, Methods). Due to the background fluorescence and scattering we observed only overall fluorescence changes in the entire field of view, which included many neurons in n=2 slices. We chose neural stimulation parameters similar to those commonly used for deep brain stimulation (So et al. 2017): 100 biphasic pulses at 200 Hz with each phase lasting 175μs. When the magnetic field was on we observed a corresponding increase in fluorescence in n=23 recordings in neocortical layer 5 consistent with activity-mediated calcium increases. Following bath application of tetrodotoxin (TTX, 500 nM) fluorescence increases were completely blocked in n=9 recordings confirming that ME evoked activity was dependent on action potentials in nearby neurons.

### ME Neural Stimulation in Freely Moving Rats Provides Therapeutic Benefit

A major advantage of our ME stimulators is the fact that remote activation enables experiments with freely behaving animals. As a proof-of-principle we adapted our biphasic stimulator for deep brain stimulation (DBS) in freely moving rats (Fig. 4). To test ME stimulator efficacy, we used a previously reported protocol to test DBS in hemi-parkinsonian rats (Summerson et al. 2014). In these experiments rats are injected with 6-OHDA in the left medial forebrain bundle (MFB) to create a unilateral lesion of the substantia nigra pars compacta (SNc). The animals are then placed in a 30 cm diameter circular enclosure. Following a dose of methamphetamine, the hemi-parkinsonian rats have been shown to rotate ipsilateral to the injection (e.g. left for injection into the left MFB). During these rotations, the rat primarily moves using its contralateral (right) forepaw, rarely placing the ipsilateral (left) forepaw onto the ground. When a biphasic stimulus is applied at 200 Hz in the sub-thalamic nucleus (STN) using a tethered electrode array stimulator, rats typically stop turning to the left and exhibit more normal behavior such as moving with both forepaws, maintaining a steady orientation, or turning to the contralateral side (So et al. 2017).

**Figure 4.**
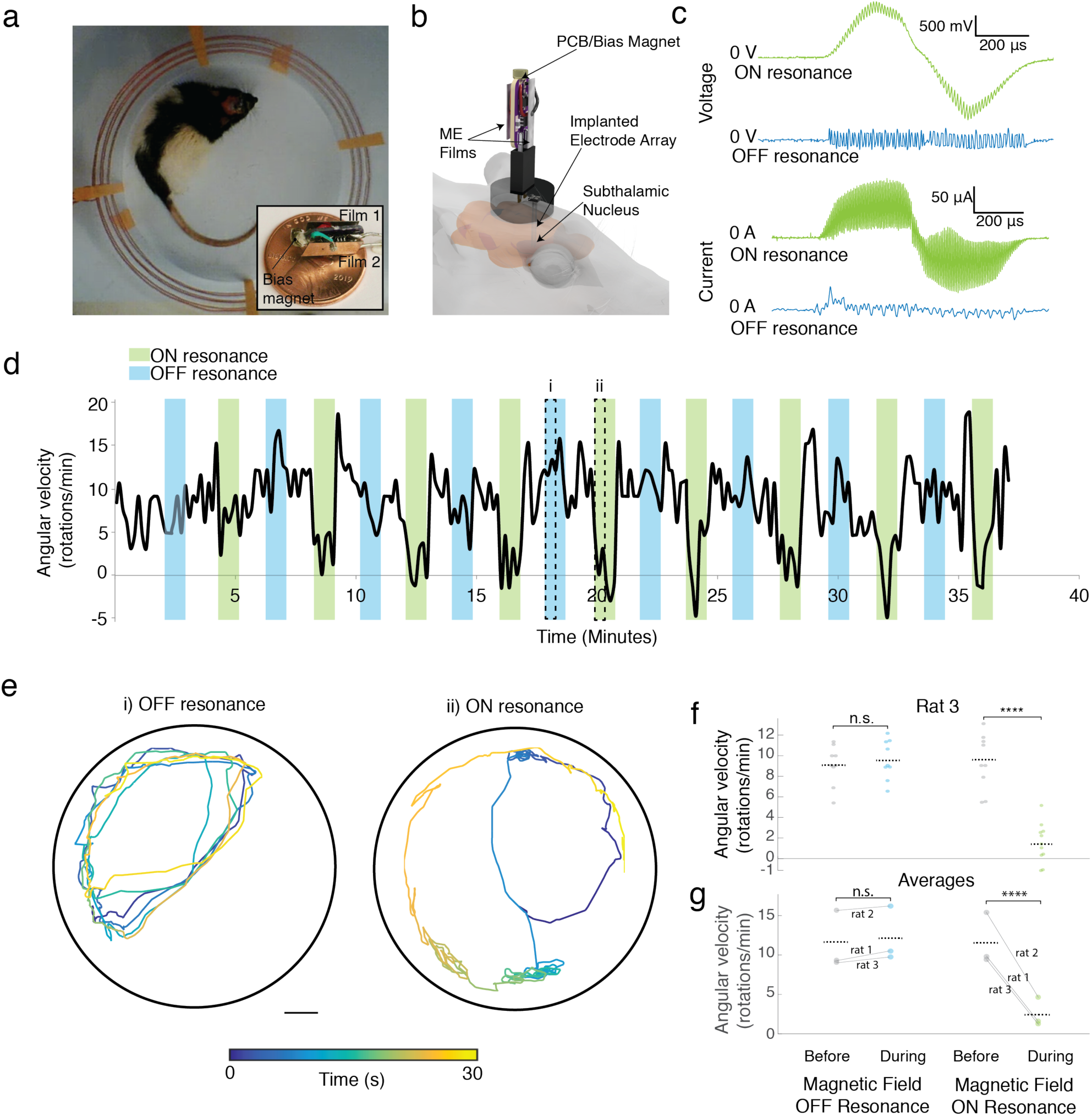
Effective DBS in a freely moving rat using a wireless ME stimulator. (a) Experimental setup showing rat in a circular enclosure wrapped with magnet wire. Inset shows a biphasic ME stimulator on a one cent coin (b) Schematic of the biphasic ME stimulator attached to the electrode array that is implanted into the STN (c) Measured voltage generated by the ME device and the current applied to the brain on resonance (green) and off resonance (blue). Approximately 100 μA biphasic stimulation is applied only when then the magnetic field frequency matches the resonance condition. (d) Angular velocity of the hemi-Parkinsonian rat over a 40 minute DBS trial with intervals of resonant and non-resonant stimulation shows that rotations are reduced only when the stimulator is activated by a resonant magnetic field (e) Typical trajectories show the location of the animal’s head over two 30-second intervals denoted in c (scale bar = 5cm) (f) Average angular velocity of the rat during the 30 seconds before stimulation and the first 30 seconds of stimulation for each interval during the 40-min experiment shows a clear reduction in angular velocity only when the ME film is activated on resonance (**** P = 4×10^−7^, n.s.=not significant P=0.70, paired t-test) (g) Average angular velocities for n=3 rats shows repeatable results across multiple animals (**** P = 2.8×10^−18^, n.s.=not significant P=0.11, paired t-test)

To create a wireless, biphasic ME stimulator for freely moving animals we added a small permanent magnet to the ME stimulator to generate a bias field, and wrapped the behavioral chamber with 18 AWG copper wire to create a solenoid (Fig. 4a, S5). By integrating the small permanent magnet (< 0.25g) into the ME stimulator, we could ensure that the bias field was constantly aligned with ME films as the animal moved within the enclosure. We could also ensure that the positive and negative stimuli had equal amplitudes by independently adjusting the distance between each film and the permanent magnet. This ME stimulator was then connected to a commercial electrode array (Microprobes) implanted in the STN (Fig 4b, see Methods). We ensured that the stimulation voltage and current were within the safe and therapeutic range by measuring the output of the ME stimulator connected to an equivalent circuit model of the brain (Fig. 4c, see Methods). Specifically, we observed peak voltages of approximately +/-1.5 V and peak currents of approximately +/- 100 μA for 400 us at approximately a 50% duty cycle (200 μs of overall current per phase), which is within the effective stimulation range reported for conventional wired stimulators (Summerson et al. 2014). When we tune the magnetic field frequency off resonance we observe almost no generated voltage or current (Fig. 4c).

We then tested the wireless version of our biphasic ME stimulator mounted to the head of a freely behaving rat and found that ME stimulation showed efficacy comparable to previously reported wired DBS stimulators (Fig. 4). With a magnetic field applied at resonance, we found that one-minute periods of 200 Hz biphasic pulses resulted in a significant decrease in the animal’s rotation rate (Fig. 4d green intervals). This decreased rotation was not observed when the magnetic stimulus frequency was tuned off resonance (Fig. 4d blue intervals). Plots of the head trajectories show that the pathological ipsilateral rotations observed during off-resonant magnetic field stimulation are not present when the ME stimulator is active during resonant magnetic field stimulation where the rotations are either not present or contralateral as expected for successful stimulation (Fig. 4e, Supplemental Video 2,3, Methods). When averaged over all trials, average rotation rate during the first half of stimulation fell to a statistically significant 1.4 rotations per minute (rpm), compared to 9.4 rpm in the absence of stimulation, or 10.6 rpm during off-resonant stimulation (paired t-test, Fig. 4f). We further demonstrated the repeatability of this stimulator by repeating this stimulation protocol on two other rats and found similar results (Fig. 4g).

With a weight of 0.5 g, the ME stimulators described here are miniature, magnetic, high frequency stimulators. Furthermore, by changing the frequency and timing of the external drive coils we can generate a variety of stimulation patterns throughout the therapeutic window of 100-200 Hz with applications to other disease models.

### Millimeter-sized ME devices enable fully implanted biphasic stimulation in freely behaving rodents

By miniaturizing the components of our ME stimulator, we created a fully implanted version of our biphasic stimulators (Fig. 5). This biphasic device is composed of two mm-sized ME films (4.3 mm × 2 mm, and 5.4 mm × 2 mm in area) connected with the same circuit elements described previously (Fig 5b) and wired to a stereotrode. We packaged the device using a 3-D printed plastic shell coated in epoxy in order to protect the films and circuit from the surrounding biological environment (Fig. S5b). This miniaturized design enabled the films and circuitry to be placed on the skull of the rat with the skin sutured up over the implant (Fig 5a,e), which could help prevent issues arising from percutaneous leads.

**Figure 5.**
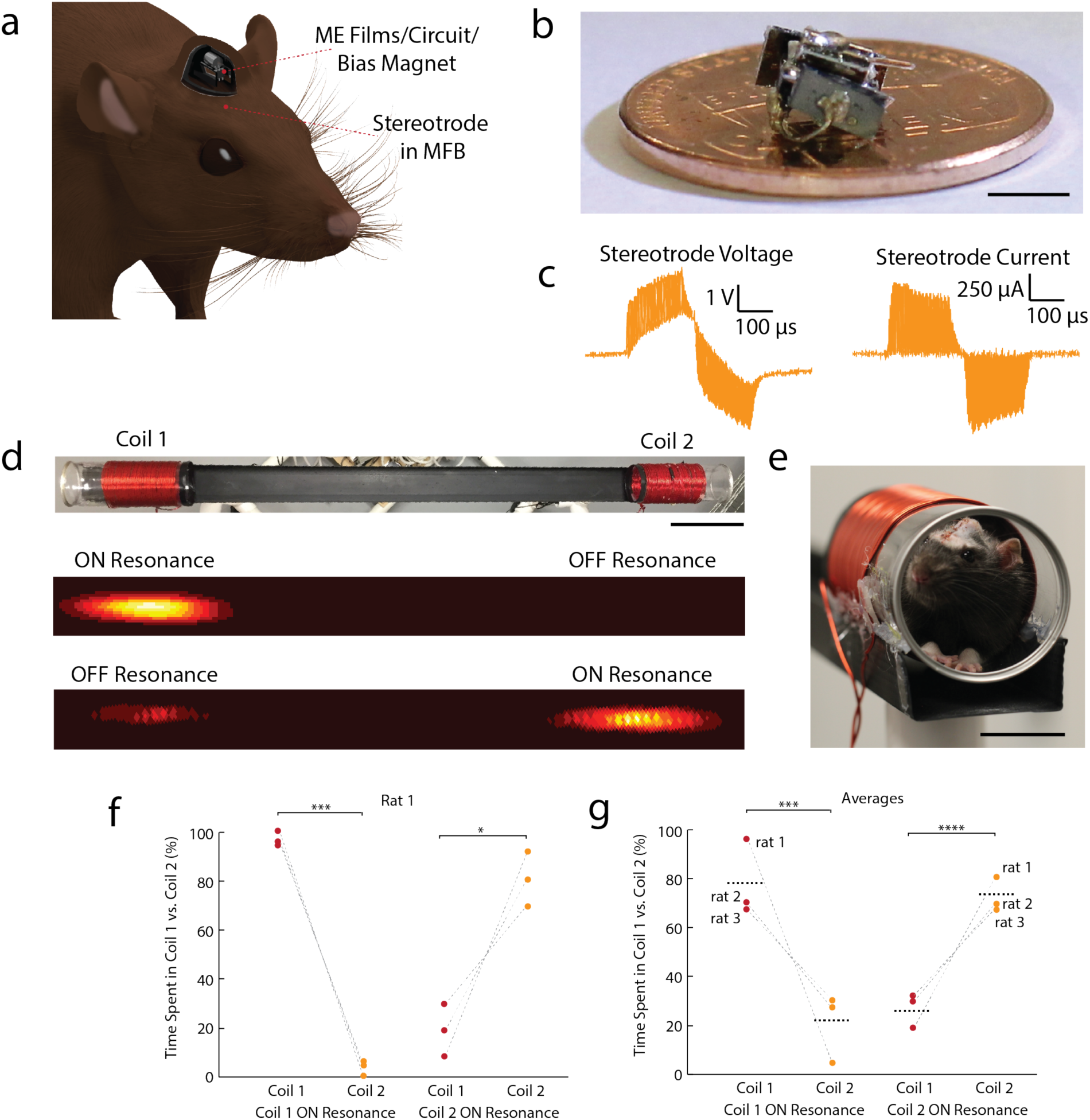
Fully implanted ME device stimulates place preference in freely moving rats. (a) Schematic of the device implanted under the skin of a rat with stereotrode implanted into the MFB (b) Photo of inner circuit and ME films used in the implant scale bar=5 mm (c) Representative voltage and current waveforms used for stimulation (d) Experimental setup showing the linear track and coil 1 and 2 and representative heat maps for two individual trials showing a location preference for the ON resonance coil demonstrating that we could change the preference by altering the magnetic field frequencies in each coil (scale bar=10 cm) (e) side view of the experimental setup showing a rat in the coil with the skin sutured up over the implant scale bar=3cm(f) Preference results from one rat over six trials (***P = 9×10^−4^, *P = 0.038) and (g) Average results for n=3 rats shows repeatability across subjects (***P = 2.7×10^−4^, ****P = 3.6×10^−5^)

To demonstrate that this fully implanted ME device effectively stimulates neural activity in a freely moving rodent, we performed a place-preference experiment. Because this experiment requires higher stimulation currents than the DBS experiment we replaced the PVDF piezoelectric layer with PZT (Fig 5c). We then implanted the device connected to a stereotrode surgically placed in the medial forebrain bundle (MFB), which is part of the reward pathway commonly used to drive behavior (Olds & Milner 1954). One to three days following surgery we placed each rat (n = 3) on a linear track with two custom-designed double resonant coils at either end of the track. For each experiment the coil at one end for the track was ON resonance for both of the implanted films and coil at the other end of the track was OFF resonance. To test for any confounding effects of the magnetic field, we ensured that each coil produced the same field strength of 1.5 mT (Fig 5d, S5a). When the rat’s head was inside the ON resonant coil, the device was activated, stimulating dopaminergic neurons in the MFB and inducing a location preference for that coil (Fig 5d, Supplementary Video 4). We quantified this effect by analyzing the amount of time the rat spent in the ON resonant coil compared to the OFF resonant coil in six 10-minute trials/rat. To demonstrate that this effect was due to the ME device we performed three trials and then switched which coil was on resonance for another three trials. As expected, we found that the place preference switched to match which coil was on resonance with our implant (Fig 5f,g, S5c,d). In each case we see a significant preference for the ON resonant coil, which confirmed that this fully-implanted biphasic device is an effective neural stimulator.

## Discussion

Here we have demonstrated a fully implanted miniature, magnetic neural stimulator that 1) enables individually addressable miniature stimulators deep within a human skull phantom and 2) operates in the therapeutic band (100-200 Hz) in freely moving animals; however, the advantages of ME materials extend beyond these proof-of-principle demonstrations.

As we demonstrated, the electric field from a ME film can directly stimulate surrounding cells *in vitro*. This means that ME materials have the potential to enable miniature neural stimulators that can be implanted deep in the brain of large animals or humans and activated externally with a small electromagnet. As shown here, rice-sized films can be selectively activated based on unique resonant frequencies. Additional miniaturization is not expected to reduce the voltage produced by these films since the voltage depends on the thickness of the piezoelectric field and not the film length (Fig. S1k), suggesting that even smaller films could serve as effective stimulators. Furthermore, in the future the rectifying diode and/or LED could be fabricated directly onto the ME film using thin film lithography techniques, which would enable miniature materials-based ME stimulators.

External ME stimulators such as the one described in the *in vivo* rat experiment could have an immediate impact on the study of DBS therapies using rodent disease models. Because the ME stimulator is compatible with commercial implanted electrodes, and the magnetic stimulators can be adapted to a number of standard behavioral experiments or animal enclosures, our ME stimulators could readily replace the wired DBS stimulators currently in use. As a result, new experiments can be developed to probe the effects of chronic and continuous DBS or DBS in social contexts where wired DBS stimulators would be impracticable.

Fully implanted ME stimulators have further advantages of avoiding routes for infection and allow for even more freedom of movement and social interaction between animals. Devices such as the ones used here could easily be adapted to target different brain areas or peripheral nerves using commercially available or custom designed electrodes.

Our preliminary lifetime testing showed good performance under physiological conditions, but more work is needed to develop packaging solutions for chronic *in vivo* applications. Specifically, we found that these miniaturized ME films maintained their functionality during a 14-day soak test in 37°C saline and when implanted in an agar tissue phantom. When encapsulated with polyimide, ME films in saline powered by a magnetic field showed no decrease in film voltage over 14 days (Fig S6e). Additionally, when we placed the polyimide-coated films into an agarose gel that closely matches the mechanical properties of brain tissue we found only a 20% decrease in voltage as a result of this mechanical dampening (Fig S6f). Because these films can produce in excess of 30 V, we expect these films to function under biological conditions. To avoid this dampening one could design packaging solutions that reduce the mechanical coupling between the film and the tissue. Future testing, including immunohistochemistry, will be needed to assess the foreign body response and develop packaging that limits this response and is stable for chronic use.

For applications in the brain or peripheral nervous system that require chronic high-frequency stimulation it will be important to measure the internal device losses due to heat and pressure waves and how these losses could affect the tissue. As a preliminary experiment, we measured the temperature of the film following 5 minutes of pulsed activation (ME Film = 6 V_pp_ and 20% duty cycle at 100 Hz) and observed no increase in the film temperature (Fig. S6c-d). Should pressure waves pose a hazard, one could develop packaging solutions that minimized the mechanical coupling between the film and the tissue.

The overall volume of the fully implanted stimulator, while small compared to conventional stimulators, remains relatively large compared to other novel implanted technologies (Table 1). However, the size of the actual ME power source is quite small (2-4 mm^3^, Table 1) while the bulk of the implant is packaging, bias magnet, and off-the-shelf circuit elements. Further substantial miniaturization is possible with improved layouts and packaging as well as with custom miniature integrated circuits. The magnetic field system could also be updated so that the bias field is included in the electromagnet setup, which would remove the need for a small on-chip permanent magnet. The realization of all this, together with further deceases in the ME film size could lead to sub-mm sized implantable devices. One potential challenge with future free-floating devices for clinical applications is that they could migrate over time from the target tissue. Indeed future work should address this issue and explore methods to anchor or tether the devices using biocompatible adhesives (Mahdavi et al. 2008), or mechanical anchors like nerve cuffs (Piech et al. 2020).

The arena size used in the rotation experiment is comparable to other experiments using RF powered devices, and future work should focus on exploring larger arenas and the associated engineering challenges. Calculations of the magnetic field strengths suggest that we can reconfigure the drive coils for a number of behavioral experiments by placing coils beneath the floor of an animal enclosure. Finite element simulations and measurements show that even at distance 4-5 cm above a drive coil, ME films generate sufficient voltage for some low levels of stimulation (Fig. S6a,b). This distance could be further improved by optimizing the geometry of the coils or increasing the power of the magnetic field.

The proof-of-concept experiments here are intended to show that ME materials provide an effective wireless power solution for miniature neural stimulators, but their performance and application space can be greatly expanded by adding application specific integrated circuits (ASICs). With these more complex circuits one could create biphasic stimulation using a single ME film or generate wirelessly programmable stimulation at various specified voltage levels (Yu et al. 2020).

Additionally, one can imagine networks of devices that can be individually addressed using wireless network protocols implemented in the ASICs. This ability to ensure safe and effective stimulation with integrated circuits will likely be required for clinical translation of this technology.

We also foresee applications for ME materials as a wireless power technology for more complex implantable bioelectronic devices. For example, the demonstrated ability of ME films to power LEDs implies that ME materials could power implantable optogenetic stimulators, small integrated circuits for physiological monitoring, or transmit data out for closed-loop bioelectronic devices.

To fully realize these new bioelectronic devices based on ME materials, work is needed to improve ME materials and fabrication processes to reliably produce high-quality miniature ME films, and encapsulate them for chronic use. For wearable technologies, it is also necessary to further miniaturize magnetic field generators so that they can be battery powered and comfortably worn. These advances must also be accompanied by in vivo testing to show safety and efficiency for chronic use.

With the benefits of ME-based bioelectronics also come limitations. The need for magnetic materials may limit the magnetic imaging compatibility of some devices. Compared to ultrasound and RF wireless power that rely on propagating waves, our ME devices are powered by near-field magnetic fields. As a result, the depth that we can effectively power ME devices depends size of the transmitter. On other hand, the magnetic fields used here show negligible absorption by the tissue allowing us to increase the power in the transmitter and remain well below the safety limits. Together, these considerations provide design tradeoffs when developing a system for miniature bioelectronic implants, where the constraints on the size of the transmitter, need to transmit through air, and total power needed at the device may lead one to choose one wireless power solution over another. Additionally, bidirectional communication based on ME effects may be difficult since the magnetic fields do not radiate like ultrasound or electromagnetic waves.

Overall, ME materials have the potential to fill a key need for wireless power delivery to miniature neural stimulators and other bioelectronic devices where the major challenge is transferring energy over distances of several centimeters without heating the tissue or suffering loss at interfaces between tissue, bone, and air.

## Acknowledgments

We would like to acknowledge our attending veterinarian at Rice University, Dr. Elysse Orchard, for help with the surgical procedures to ensure our devices stayed fully enclosed underneath the skin.

## Author Contributions

## Declaration of Interests

The authors declare no competing interests.

## STAR Methods

### Resource Availability

#### Lead Contact

Further information and requests for resources and reagents should be directed to the Lead Contact, Jacob Robinson.

#### Materials Availability

This study did not generate new unique reagents.

#### Data and Code Availability

The arduino teensy code used to drive the magnetic fields can be found at: https://github.com/RobinsonLab-Rice/MagElecCoilDriver

### Experimental Models and Subject Details

#### Cell Lines

HEK cells expressing sodium channel Na_1.3_ and potassium channel K_2.1_ were obtained from the lab of Dr. Adam Cohen (Harvard). Cells were cultured and maintained at 37°C in cell culture media (DMEM/f12 with strep, pen, geneticin, and puromycin)

#### Animals

##### Brain slice experiments

All experiments were performed in accordance with NIH guidelines and approved by the University of Texas Health Science Center at Houston (UTHealth) animal welfare committee. For the electrophysiology, we used 16-day-old C57BL6/J mice (JAX #000664). For the GCaMP imaging we used 40 day old GAD2-GCaMP3 mice, generated by crossing GAD2-Cre (JAX # 10802) with flox-GCaMP3 (JAX # 14538) animals.

##### In vivo experiments

All experiments were approved by the Rice University Institutional Animal Care and Use Committee’s guidelines and adhered to the National Institute of Health guidelines. Six adult male Long-Evans rats aged 4-7 months and weighing 500-800 grams from Charles River Laboratories were used for this study. Rats were housed in pairs prior to surgery and randomly chosen for implantation. Post-implantation rats were individually housed. At all times animals were kept on a 12 hour light-dark cycle.

### Methods Details

#### Film Fabrication

To fabricate ME films, we used Metglas SA1 alloy (Metglas Inc) for the magnetostrictive layer and polyvinylidenefluoride “PVDF” (precision acoustics) or lead Zirconate titanate “PZT” (Piezo Systems) for the piezoelectric layer. The PVDF films used for these experiments were pre-stretched and poled by the manufacturer. The two layers were bonded together using an epoxy capable of transferring the mechanical stress between the two layers (Hardman double bubble red epoxy). Prior to bonding the two layers together, we sputtered a thin layer of platinum (<100 nm) as a top electrode on the PVDF. Both the Metglas and PVDF were plasma cleaned using O_2_ plasma for five minutes before epoxying. After the epoxy set, the films were cut into the desired rectangular shape using scissors, taking care to cut the long axis of the film along the stretching direction of the PVDF. We then attached wires using conductive epoxy to either side of the films in order to measure the electrical capabilities of the film. We found that attaching wires in the center dramatically increased the resonant voltage. However for convenience, the wires were attached near the ends of the films during the in vitro experiments. In many cases we also attached additional electronic components such as diodes or LEDs to the wires attached to the films as noted in the appropriate sections in the main text. Finally the devices were coated in 5-10 μm of parylene-C (Labcoater 2). Initially this coating was used to electrically insulate and protect the devices during in vitro experiments, but we also found that the encapsulation increases the resonant voltage, which could be due to increased mechanical coupling from the encapsulation.

#### Bench Top Electrolysis Tests

The stimulator shown in Fig 4a was wired to a stereotrode immersed in saline under a microscope in order to observe the formation of bubbles from electrolysis at the tips. During monophasic stimulation we used only one resonant frequency and during biphasic stimulation we used two frequencies as demonstrated above. In each case the pulse time was a 400 μs/phase. We determined the limit of stimulation time as when the first bubble began to appear at the tips of the electrode and repeated each data point 4 times.

#### Magnetic Field Generation (Fig. S1)

Each magnetic field generator consists of two major components, 1) Magnetic coils used for the alternating magnetic field (described in the main text and optimized for each experiment) and 2) Electronic drivers to control voltage and timing of the alternating current in the coils (the same for all experiments).

To maintain simplicity, efficiency, and low cost the coils were driven with full H-Bridge style switching circuits. The drivers are designed to deliver high currents to the drive coils in the form of bi-phasic pulse trains. This reduces the cost and complexity of the driver itself, as well as the power supply and control circuitry when compared to arbitrary function generators. The design also has potential for improved operational efficiency through impedance matching with the drive coils. Furthermore, it is also possible to regulate power delivered to the drive coils on the fly by adjusting the duty cycle of the current pulses, allowing power being delivered to the ME film to be easily controlled digitally while maintaining the resonant carrier frequency. The output carrier and pulse frequencies of the magnetic field are generated using a TeensyLC board and custom Arduino code to generate the specific pulse timings to deliver controlled ME stimulation (Fig. S1e-h).

These coils and drivers can be combined in different ways to generate the appropriate field for a given experiment. For example, the setup used to generate the alternating field in the in vivo rotation experiments consisted of four sets of coils each with five turns powered by one driver with all four drivers synced to the same output signal. In this way we can generate sufficient power to generate a mT-scale magnetic fields over the whole behavioral area (Fig. S4a-b).

#### Resonant Coil Design

In order to generate sufficient field strengths of >1mT at the resonant frequencies for the miniature devices (300-400 kHz) we developed a custom resonant coil system as shown in Fig S5. This system had two separate resonant frequencies to be able to selectively active each film on the miniature device. The first higher resonance is determined by a single capacitor C_1_ in series with the behavioral coil while the self-resonance from a second inductor acts as a low pass filter to prevent a second capacitor C_2_ from affecting the system. The second lower series resonance is determined from adding C_1_ and C_2_ in parallel as the inductor no longer filters out the lower resonant frequency.

#### In Vitro HEK Experiments

For experiments performed on coverslips, HEK cells expressing sodium channel Na_1.3_ and potassium channel K_2.1_ were grown on 12 mm poly-l-lysine coated coverslips to approximately 30% confluency. The cells were then transfected with the genetically encoded voltage indicator ArcLight using Lipofectamine (Invitrogen) following manufacturer’s recommendations. Two to three days after transfection the coverslips were inverted onto ME films for testing. Preparation of GFP controls followed the same procedure with the exception of replacing the ArcLight vector (AddGene) with a GFP expression vector (AddGene). For experiments performed with cells grown on the films, HEK cells transfected with ArcLight were placed onto parylene coated poly-l-lysine treated films. The films were placed in cellular media overnight and tested the following day.

ArcLight and GFP were excited at 460 nm with an LED light source. Fluorescence images were collected at 33 fps using a CCD camera. Images were analyzed using Matlab to quantify fluorescence changes in individual cells. In vitro testing was performed in extracellular buffer (ECB, in mM: NaCl 119, KCl 5, Hepes 10, CaCl_2_ 2, MgCl_2_ 1; pH 7.2; 320mOsm)

Figure S2b was obtained by growing unmodified HEK cells on a film submerged in cellular media for five days. The cells were then stained with Hoechst and Calcein-AM to label the nucleus and membrane respectively in living cells. The cells were then fixed and imaged using a confocal microscope.

#### Skull Phantom Demonstration

At the magnetic field frequencies used for this experiment bone and tissue are effectively transparent (Bottomley & Andrew 1978), so we selected a life sized skull with the size of an average human adult head as a phantom (Orient Infinity Limited). It was wrapped with 18 AWG magnet wire as shown in Fig 2. The coil consisted of four coils in parallel each wired to an individual magnetic field driver. All drivers were wired to the same input frequency signal and powered from the same power supply. The films were suspended at the center of the skull phantom. Orange LEDs (Chanzon) with a diode antiparallel were attached to the films for wireless verification of the voltage generated by the films. For visualization purposes the skull top was removed to better photograph the LED.

#### Electrophysiology

We prepared thalamocortical brain slices (400 μm) as described previously (Agmon & Connors 1991). Briefly, animals were anesthetized and decapitated, in accordance with NIH guidelines and approved by the University of Texas Health Science Center at Houston (UTHealth) animal welfare committee. Brains were removed and immediately transferred to an ice-cold sucrose-based cutting solution containing (in mM): 234 sucrose, 2.5 KCl, 1.25 NaH_2_PO_4_, 10 MgSO_4_, 0.5 CaCl_2_, 26 NaHCO_3_, and 11 glucose, saturated with 95% O2, 5% CO_2_. Slices were cut using a vibratome (Leica VT 1200S) and transferred to ACSF containing (in mM): 126 NaCl, 2.5 KCl, 1.25 NaH_2_PO_4_, 2 MgCl_2_, 2 CaCl_2_, 26 NaHCO_3_, and 10 glucose. Slices were held at 35°C for 20 min and then kept at room temperature until used for recordings. For experiments, slices were placed in a recording chamber and perfused with ACSF held at 31-34°C and containing NBQX (R&D Systems) to block AMPA receptor-mediated synaptic transmission. Whole-cell voltage-clamp recordings from neurons in the thalamic reticular nucleus (TRN) were performed using glass pipettes (3-5 MΩ) filled with a potassium based internal solution containing (in mm): 133 K-gluconate, 1 KCl, 2 MgCl_2_, 0.16 CaCl_2_, 10 HEPES, 0.5 EGTA, 2 Mg-ATP, and 0.4 Na-GTP (adjusted to 290 mOsm, pH 7.3). Antidromic action potentials were evoked by placing stereotrodes in the adjacent ventrobasal nucleus of the thalamus. Data were acquired using a Multiclamp 700B amplifier (Molecular Devices), filtered at 10 kHz, and digitized at 20 kHz with a 16-bit analog-to-digital converter (Digidata 1440A; Molecular Devices).

#### GCaMP Imaging

Brain slices were prepared closely following procedures described by Ting et al (Ting et al. 2014). Mice were deeply anesthetized with Isoflurane and perfused with ice cold NMDG-based solution consisting of (in mM): 92 NMDG, 2.5 KCl, 1.25 NaH_2_PO_4_, 10 MgSO_4_, 0.5 CaCl_2_, 30 NaHCO_3_, 20 glucose, 20 HEPES, 2 thiouera, 5 Na-Ascorbate, 3 Na-pyruvate, saturated with 95% O_2_ and 5% CO_2_., at a rate of ∼6 ml/min. Coronal brain slices (300 µm) were cut using a vibratome (Leica VT1200S), incubated for 15 min at 35 °C in NMDG-based solution, and then transferred to a chamber held at room temperature containing (in mM): 92 NaCl, 2.5 KCl, 1.25 NaH_2_PO_4_, 2 MgSO_4_, 2 CaCl_2_, 30 NaHCO_3_, 25 glucose, 20 HEPES, 2 thiouera, 5 Na-Ascorbate, 3 Na-pyruvate, saturated with 95% O_2_ and 5% CO_2_. For experiments, slices were placed into a recording chamber perfused with ACSF containing (in mM): 126 NaCl, 2.5 KCl, 1.25 NaH_2_PO_4_, 2 MgCl_2_, 2 CaCl_2_, 26 NaHCO_3_, 10 glucose), held at 32-34 °C. NBQX (10 μM) was included in the bath solution to block AMPA receptor-mediated synaptic transmission. Stereotrodes were placed in layer 5 of somatosensory (barrel) cortex. GCaMP3 expressed in GAD2 positive GABAergic neurons was excited at 460 nm with an LED light source. Fluorescence images were collected at 9.8 fps using a CCD camera attached to an Olympus BX51WI microscope. Images were analyzed using Matlab to quantify fluorescence changes in 600 x 600 μm regions around the stereotrode tips.

#### Rat Surgical Procedures

All experiments were approved by the Rice University Institutional Animal Care and Use Committee’s guidelines and adhered to the National Institute of Health guidelines. For both rotation tests and place preference animal experiments a total of six male Long-Evans rats (three per experiment weighing in the range of 500-800 grams) were anesthetized with isoflurane gas. Five percent isoflurane was used to induce anesthesia and 1.5-2.5% was used to maintain anesthetic depth. Buprenorphine (0.04mg/kg) was administered prior to ear bars as an analgesic. Following initial setup surgical methods differ for the two experimental protocols. For rotation test experiments, 5-7 skull screws were placed to anchor the electrode array. Skull screws were bound to skull with C&B Metabond (Parkell). A craniotomy was made to accommodate the microelectrode array and expose an injection site for neurotoxin. A 30-gauge needle was bent at the tip to pull away dura covering the brain. Desipramine (DMI) reconstituted in saline at a concentration of 15 mg/mL was injected IP to protect noradrenergic neurons. The dose of DMI was approximately 15 mg/kg and injected approximately 30 minutes prior to administration of neurotoxin. To induce hemiparkinsonian lesion, 8 ug of 6-hydroxydopamine (OHDA) at 2ug/uL in saline was injected at 0.2 uL/min into the medial forebrain bundle (MFB −1.2mm -- −1.25mm ML, −4mm AP and −8.1mm DV). STN stimulation was delivered via 2×2 platinum iridium microelectrode array (Microprobes) with 600 × 600 um spacing of 75 um electrodes. Each electrode had a nominal 10 kOhm impedance. For two rats electrode array was lowered to −2.6mm ML, −3.6mm AP and −8.2mm DV from bregma. For the third rat the electrode array was placed into the brain at 2.5mm ML, −3.4mm AP relative to bregma and −7.7mm DV relative to dura. The array was fixed to the skull with standard two-part dental acrylic. For place preference experiments 3-4 skull screws were placed above bregma. Skull screws were similarly bound to the skull with Metabond dental acrylic; however, the Metabond acrylic was limited to flow only above bregma. A craniotomy and duratomy above the location of the MFB (−1.8mm ML, −2.8mm AP) was made to accomodate for a 9 mm platinum iridium bipolar stereotrode from Microprobes with a nominal 10 kOhm impedance at 1 kHz. A custom 3D printed rounded/smoothened enclosure housing the electronics and connecting to the stimulating electrode was stereotactically lowered into the exposed brain to the depth of the MFB at −8.6mm --8.7mm DV relative to bregma. Kwiksil from World Precision Instruments was injected into the duratomy site built up to the base of the housing. Metabond was then applied again to the base of the 3D printed housing down to the Metabond on the skull screws. For a final securing of the implant, UVcuring Flow-IT (Pentron) was used to cover the implant and anchor it to the Metabond in order to avoid the heat generated from the curing of standard two-part dental acrylic from damaging the custom housing and electronics. Lastly, the animal skin was sutured over the implant leaving it enclosed underneath the skin. The sutures were found to be strongest and confirmed to hold for at least one month when using minimal (1 -- 2) interrupted sutures over the implant itself.

#### Hemi-Parkinsonian Experiments

Prior to stimulating each rat with the magnetoelectric stimulator, the stimulator power was estimated via a benchtop approximation of the rodent electrode impedance. Constant current stimulation of the rodent brain with an A-M Systems 4100 stimulator produced characteristic voltage waveforms that approximated a simplified parallel RC circuit. A 56 kOhm resistor, and 440 pF capacitor in parallel closely approximated the impedance characteristics of the rat brain across the stimulating electrodes. Using this circuit model, we estimated the field strengths and pulse durations necessary to produce the desired stimulation effects and confirm that the stimulation was charge balanced prior to rodent experimentation.

Prior to performing the rotation tests the rat was briefly anesthetized with 5% isoflurane gas and injected intraperitoneally (IP) with methamphetamine (0.31 ml 1.25 mg/kg) and the wireless biphasic stimulator was plugged into the implanted electrode array. After the anesthesia had worn off (about 5-10 min) the rat was placed in the cylindrical behavioral chamber. The magnetic field was applied over the whole behavioral area to the films on the device (Fig S5a).

The magnetic field was applied on resonance and off resonance for one minute at various times during the 40-minute trial. The resonant frequencies were 130 kHz and 160 kHz and the off resonant frequencies were 120 kHz and 170 kHz.

#### Rodent Tracking

Head position on the rotation task was generated using a slightly modified version of DeepLabCut (Mathis et al. 2018) to track ears, snout, and implant. A dataset totaling 286 frames from both the on and off resonance rotation tasks was hand labeled and trained for approximately 140,000 iterations.

Position during the place preference task was manually tracked via custom python scripts. The overall preference was quantified by counting the number of frames in which the animal was inside either one of the coils.

#### Place Preference Implant Design

The miniature ME films were fabricated from PZT/Metglas. The circuit components used were the same as those used in the brain slice and rotation test, however in this case there was no circuit board and the elements were soldered together directly and the films attached with conductive epoxy. The films/circuit were parylene coated for extra insulation and then a small bias magnet was attached in the orientation and position that ensured the best charge balance between the two phases. The plastic case designed to put the least stress on the skin was 3D printed in plastic and included a channel to securely hold the stereotrode and a chamber to hold the circuit/films. The circuit/films were attached to the stereotrode with conductive epoxy and a combination of Flow-It ALC (Pentron) and epoxy were used to encapsulate the entire outside of the box. Prior to implantation the implant was put through a 12 hour ethylene oxide cycle followed by a 12-24 hour degas period inside of a fume hood.

#### Place Preference Experiments

The rats were given a minimum of 24 hours to recover. We performed the experiments 1-3 days after the surgery. The rat was placed on a linear track with a coil at each end. Both coils applied a resonant alternating magnetic field, but only one was on resonance to activate the implanted device. We took a 10 minute video recording of the rat position starting from the first time the rat received stimulation from the ON resonant coil. After 10 minutes the rat was removed from the track and placed back into the home cage for a minimum of 5 minutes while the track was cleaned. In order to ensure that the rat did not develop associations with specific places in the room, we rotated the track to an arbitrary angle between each of the six trials. In total we performed 6 trials/rat. Between the third and fourth trial the system was also re-tuned to switch which coil was resonant with the device in addition to rotating the track.

### Quantification and Statistical Analysis

Error bars in Figure S1m denote +/- one standard deviation for n=∼50 data points, n refers to an individual film. We furthermore performed a Tukey’s Honest Significant Difference test on the data in Figure S1m, which indicated that the voltage produced at each different PVDF thickness is significantly different. Paired t-tests were used for the rodent tests in figures 4 and 5. Star values indicate levels of significance associated with the P-values listed in the figure captions. In figure 4f we compared n=9 data points for before and during off resonance stimulation and n=9 data points for before and during on resonance stimulation where n refers to 1 stimulation period during a trial. In figure 4g we compared n=29 for on resonance data points and n=28 for off resonance data points, where n refers to all the rotation rate data points for all rats during the specified time periods (averages for each rat shown in figure, not individual data points). In figure 5f we compared n=3 data points for each coil during a trial, where n=amount of time spent in one coil compared to the other. In figure 5g we compared n=9 data points (averages for each rat shown in figure, not individual data points) where n=individual data points for amount of time spent in one coil during a trial.

## Supplemental Text and Figures

**Supplemental Figure 1.**
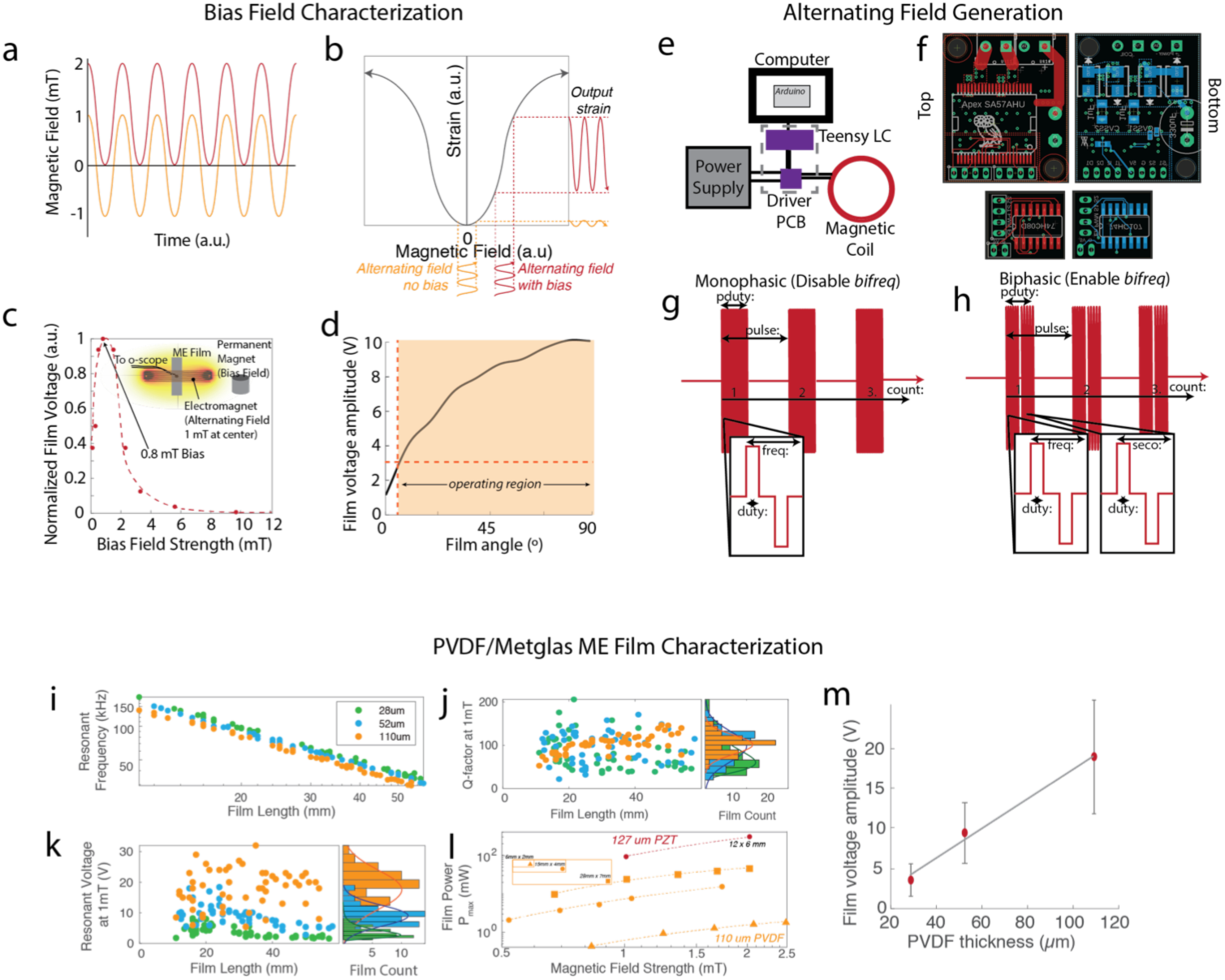
Magnetic Field and Film Characterization. (a) The magnetic field used at the location of the film in every experiment is the combination of a bias field and an alternating field (b) This maximizes the strain in the magnetoelectric material (c)The peak resonance voltage is significantly increased by a modest bias field that can be produced by a permanent magnet (d) The ideal orientation of the films is parallel with the field direction however, operation is still possible even with misalignment due to the high initial voltage (e) Schematic of the major components of the magnetic field driver. Circuit diagrams for the driver PCBs shown in (f). (g) Output waveform for monophasic stimulation and the parameters that can be controlled by the drive software (h) Output waveform for biphasic stimulation, and the parameters that can be controlled by the driver software (i) As film length decreases the resonant frequency increases but the q-factor (j) and output voltage (k) remain the same. (l) The output power depends on the film area and material, while the voltage depends on the thickness of the piezoelectric layer (m).

**Supplemental Figure 2.**
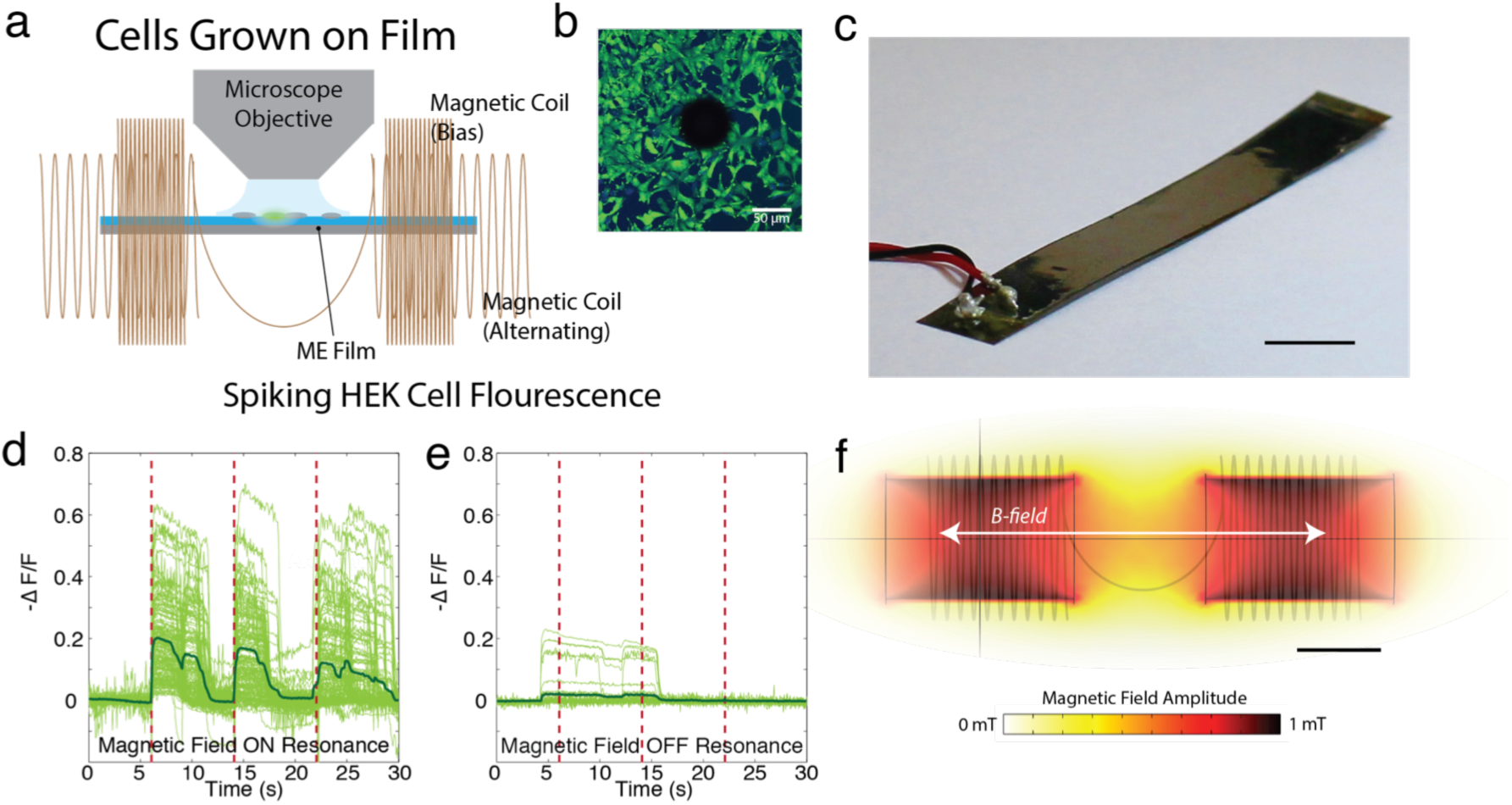
ME films *in vitro*. (a) Schematic of experimental setup for testing cells grown on ME films (b) Microscope image of fixed cells adherent to the region around a stamped hole (Hoechst/Calcein-AM, cells labeled prior to fixing) (c) sample film used for *in vitro* testing (scale bar = 4 mm) (d) ArcLight fluorescence of spiking HEK cells when magnetic field is on resonance and (e) off resonance (f) COMSOL simulation of the magnetic field strength used in this experiment (scale bar = 3 cm)

**Supplemental Figure 3.**
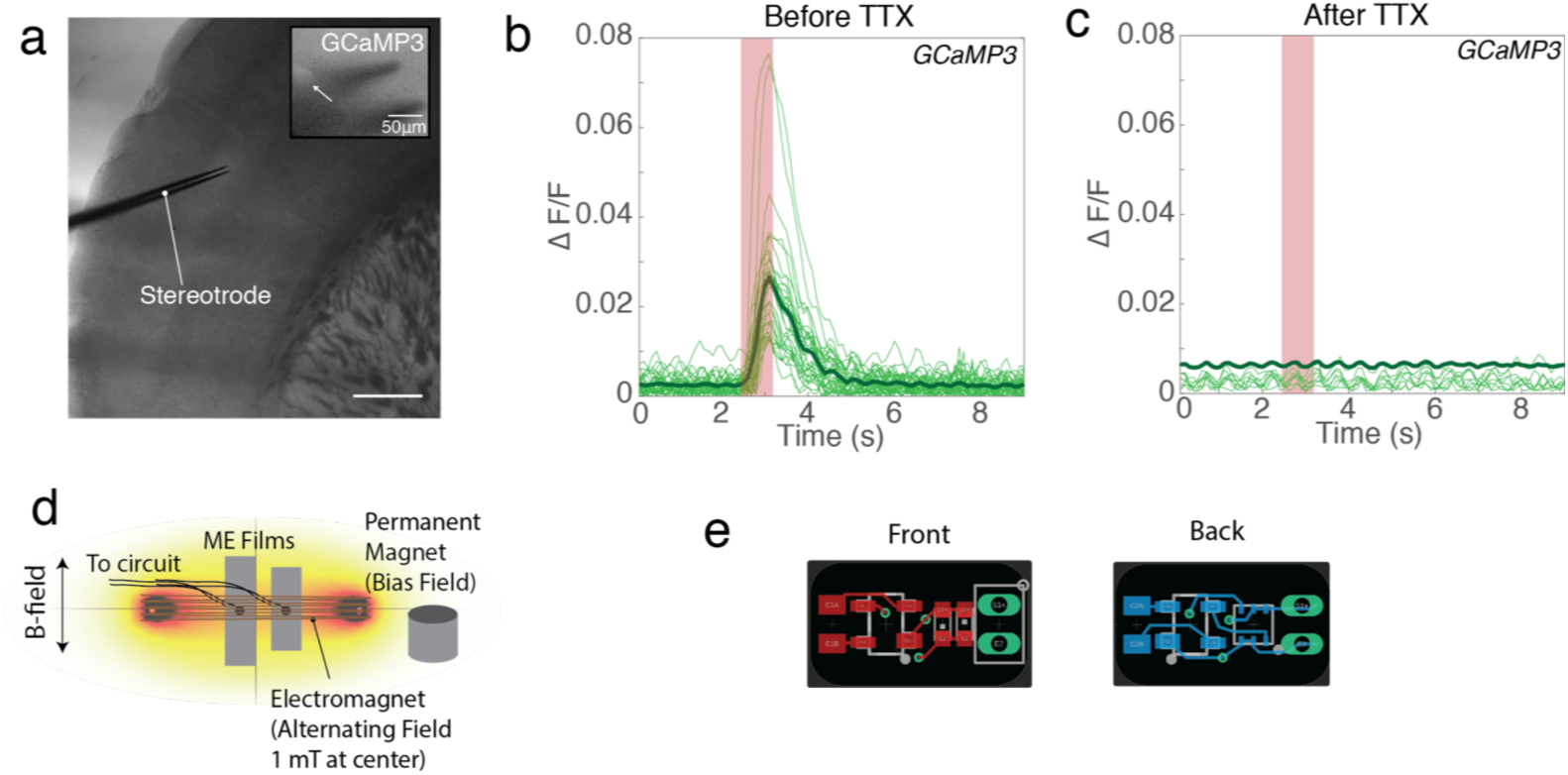
Additional Brain Slice Experiments. (a) Bright field image of stereotrode in mouse cortex (scale bar = 1 mm) with inset of GCaMP signal averaged over a 600 μm x 600 μm region around stereotrode tip. Arrow indicating a fluorescing cell body near the stereotrode (b) Time-locked increases in GCaMP signal following application of resonant magnetic field shows neural activity is induced by the ME stimulator. (c) TTX application eliminates all activity. Thin traces in (b) and (c) represent separate experiments from two different brain slices, and thick traces represent the mean of all experiments. (d) Magnetic field setup used in this setup and (e) circuit board used in this experiment

**Supplemental Figure 4.**
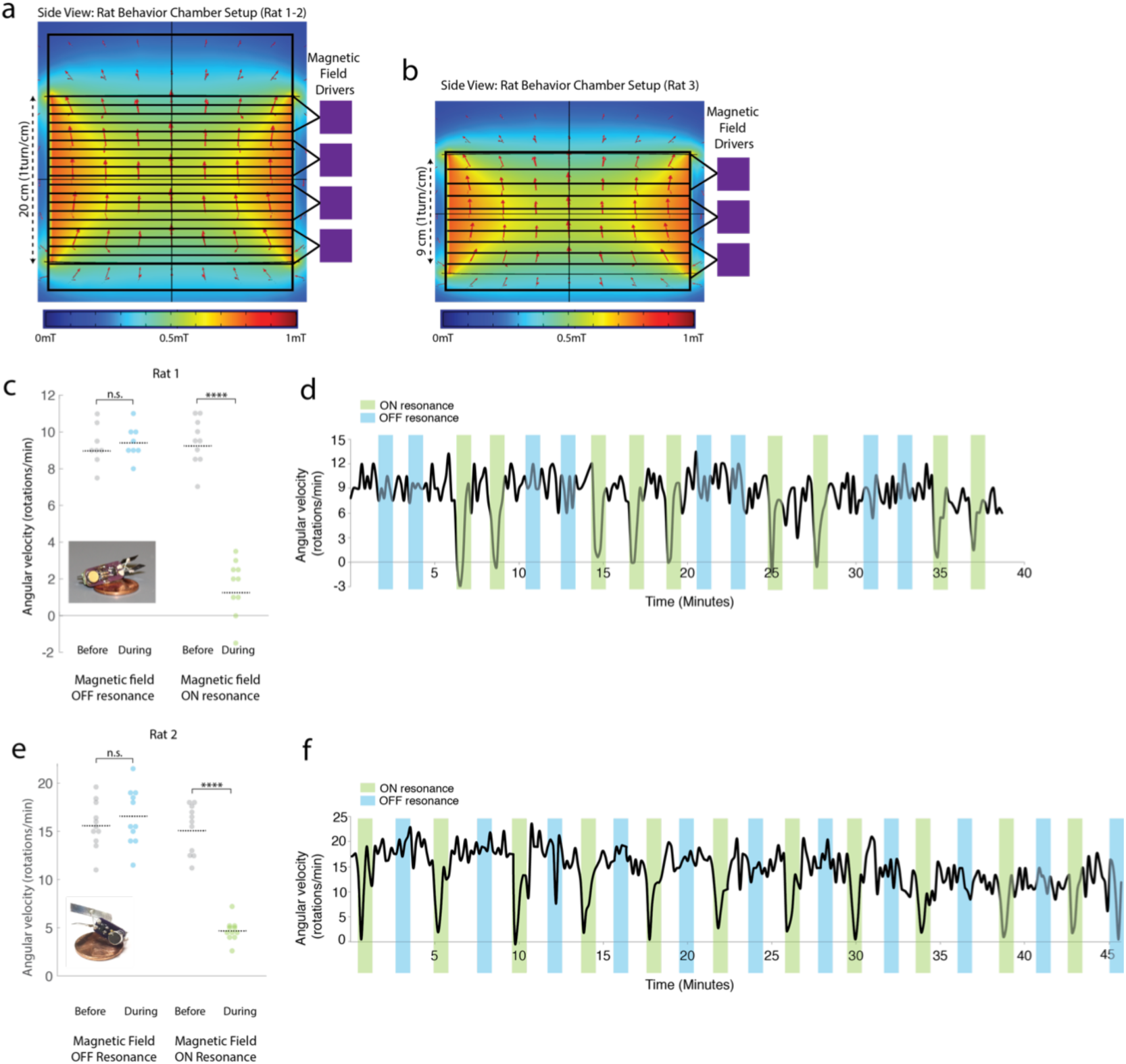

**Supplemental Figure 5.**
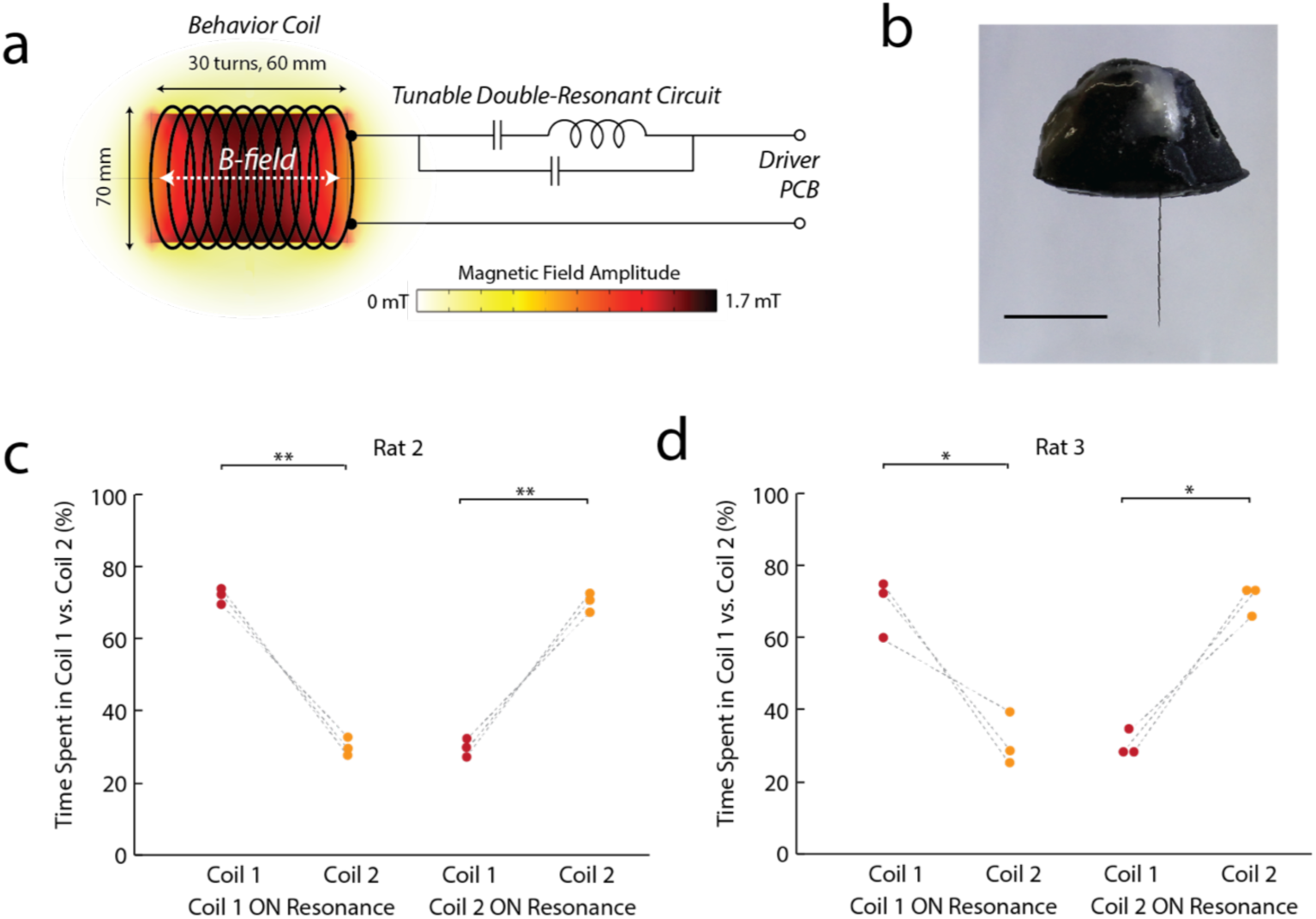
Place Preference Experiment. (a) Schematic of experimental setup showing the magnetic field circuit and magnetic field strength (b) Image of a fully enclosed ME film implant scale bar = 5 mm (c-d) Individual results of the place preference trials for rats 2 and 3

**Supplemental Figure 6.**
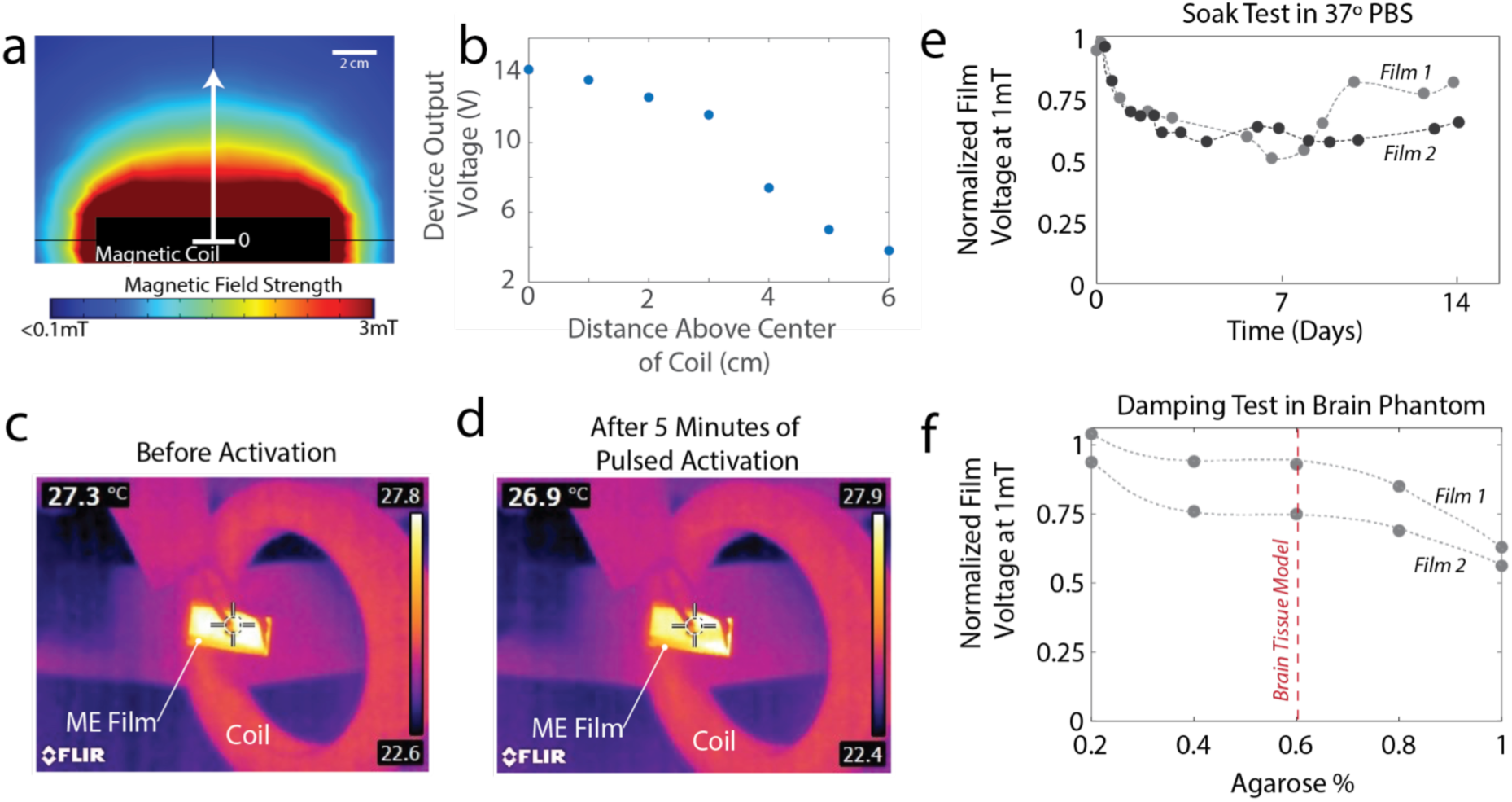
Future Considerations for ME Devices. (a) COMSOL simulation of magnetic field above a circular coil and (b) Measured device output voltage as a function of distance above the coil (c-d) Films do not heat up during five minutes of pulsed operation (e) Soak test at 37C shows that film output voltage remains relatively constant for up to 14 days of constant activation (f) Films still show usable voltage even with some damping in agarose brain phantom

## References

Agmon, A. & Connors, B.W., 1991. Thalamocortical responses of mouse somatosensory (barrel) cortexin vitro. Neuroscience, 41(2–3), pp.365–379. Available at: https://www.sciencedirect.com/science/article/pii/030645229190333J [Accessed February 26, 2020].

Agrawal, D.R. et al., 2017. Conformal phased surfaces for wireless powering of bioelectronic microdevices. Nature Biomedical Engineering, 1(3), pp.1–16. Available at: http://www.nature.com/authors/editorial_policies/license.html#terms.

Alonso, P. et al., 2015. Deep brain stimulation for obsessive-compulsive disorder: A meta-analysis of treatment outcome and predictors of response. PLoS ONE, 10(7), pp.1–16.

Amar, A. Ben, Kouki, A.B. & Cao, H., 2015. Power approaches for implantable medical devices. Sensors (Switzerland), 15(11), pp.28889–28914.

Baizabal-Carvallo, J.F. & Alonso-Juarez, M., 2016. Low-frequency deep brain stimulation for movement disorders. Parkinsonism and Related Disorders, 31, pp.14–22.

Bewernick, B.H. et al., 2010. Nucleus Accumbens Deep Brain Stimulation Decreases Ratings of Depression and Anxiety in Treatment-Resistant Depression. Biological Psychiatry, 67(2), pp.110–116.

Bottomley, P.A. & Andrew, E.R., 1978. RF magnetic field penetration, phase shift and power dissipation in biological tissue: Implications for NMR imaging. Physics in Medicine and Biology, 23(4), pp.630–643.

Carrey, J. et al., 2015. Simple models for dynamic hysteresis loop calculations of magnetic single-domain Simple models for dynamic hysteresis loop calculations of magnetic single-domain nanoparticles : Application to magnetic hyperthermia optimization. Journal of Applied Physics, 83921(2011).

Chen, R. et al., 2015. Wireless magnetothermal deep brain stimulation. Science (New York, N.Y.), 347(6229), pp.1477–80. Available at: http://www.ncbi.nlm.nih.gov/pubmed/25765068.

Fotopoulou, K. & Flynn, B.W., 2011. Wireless Power Transfer in Loosely Coupled Links : Coil Misalignment Model. IEEE Transactions on Magnetics, 47(2), pp.416–430.

Freeman, D.K. et al., 2017. A sub-millimeter, inductively powered neural stimulator. Frontiers in Neuroscience, 11(NOV), pp.1–12.

Guduru, R. et al., 2015. Magnetoelectric “spin” on stimulating the brain. Nanomedicine, 10(13), pp.2051–2061. Available at: http://www.futuremedicine.com/doi/10.2217/nnm.15.52.

Hargreaves, D.G., Drew, S.J. & Eckersley, R., 2004. Kirschner Wire Pin Tract Infection Rates: A randomized controlled trial between percutaneous and buried wires. The Journal of Hand Surgery: British & European Volume, 29(4), pp.374–376. Available at: http://www.sciencedirect.com/science/article/pii/S0266768104000841.

De Hemptinne, C. et al., 2015. Therapeutic deep brain stimulation reduces cortical phase-amplitude coupling in Parkinson’s disease. Nature Neuroscience, 18(5), pp.779–786.

Ho, J.S. et al., 2015. Self-Tracking Energy Transfer for Neural Stimulation in Untethered Mice. Physical Review Applied, 24001, pp.1–6.

International, I. & Safety, E., 2006. IEEE Standard for Safety Levels with Respect to Human Exposure to Radio Frequency Electromagnetic Fields, 3 kHz to 300 GHz,

Jin, L. et al., 2012. Single Action Potentials and Subthreshold Electrical Events Imaged in Neurons with a Fluorescent Protein Voltage Probe. Neuron, 75(5), pp.779–785. Available at: http://dx.doi.org/10.1016/j.neuron.2012.06.040.

Johnson, B.C. et al., 2018. StimDust : A 6. 5mm 3, Wireless Ultrasonic Peripheral Nerve Stimulator with 82 % Peak Chip Efficiency. 2018 IEEE Custom Integrated Circuits Conference.

Kesar, T.M. et al., 2010. Novel Patterns of Functional Electrical Stimulation Have an Immediate Effect on Dorsiflexor Muscle Function During Gait for People Poststroke. Physical Therapy, 90(1), pp.55–66. Available at: https://academic.oup.com/ptj/article-lookup/doi/10.2522/ptj.20090140.

Kulkarni, a. et al., 2014. Giant magnetoelectric effect at low frequencies in polymer-based thin film composites. Applied Physics Letters, 104(2), pp.0–5.

Maeng, L.Y. et al., 2019. Behavioral validation of a wireless low-power neurostimulation technology in a conditioned place preference task. Journal of Neural Engineering, 16(2), p.26022. Available at: http://stacks.iop.org/1741-2552/16/i=2/a=026022?key=crossref.00395b6610fa6ce2e4c1867ce3c4ce86.

Mahdavi, A. et al., 2008. A biodegradable and biocompatible gecko-inspired tissue adhesive. Proceedings of the National Academy of Sciences of the United States of America, 105(7), pp.2307–2312.

Markwardt, N.T., Stokol, J. & Rennaker, R.L., 2013. Sub-meninges implantation reduces immune response to neural implants. Journal of Neuroscience Methods, 214(2), pp.119–125. Available at: http://dx.doi.org/10.1016/j.jneumeth.2013.01.020.

Mathis, A. et al., 2018. DeepLabCut: markerless pose estimation of user-defined body parts with deep learning. Nature Neuroscience, 21(September). Available at: http://dx.doi.org/10.1038/s41593-018-0209-y.

Merrill, D.R., Bikson, M. & Jefferys, J.G.R., 2005. Electrical stimulation of excitable tissue: Design of efficacious and safe protocols. Journal of Neuroscience Methods, 141(2), pp.171–198.

Montgomery, K.L. et al., 2015. Wirelessly powered, fully internal optogenetics for brain, spinal and peripheral circuits in mice. Nature Methods, 12(10), pp.969–974. Available at: http://dx.doi.org/10.1038/nmeth.3536.

Mulpuru, S. et al., 2017. Cardiac Pacemakers : Function, Troubleshooting, and Management. Journal of the American College of Cardiology, 69(2).

Munshi, R. et al., 2017. Magnetothermal genetic deep brain stimulation of motor behaviors in awake, freely moving mice. eLife, 6, pp.1–26.

Nan, T. et al., 2017. Acoustically actuated ultra-compact NEMS magnetoelectric antennas. Nature Communications, 8(1), pp.1–7. Available at: http://dx.doi.org/10.1038/s41467-017-00343-8.

Nurmikko, A. V, 2018. Approaches to large scale neural recording by chronic implants for mobile BCIs. In 2018 6th International Conference on Brain-Computer Interface (BCI). pp. 1–2.

O’Handley, R.C. et al., 2008. Improved Wireless, Transcutaneous Power Transmission for <emphasis emphasistype=“italic”>In Vivo</emphasis> Applications. IEEE Sensors Journal, 8(1), pp.57–62.

Olds, J. & Milner, P., 1954. Positive reinforcement produced by electrical stimulation of septal area and other regions of rat brain. Journal of comparative and physiological psychology, 47, pp.419–427.

Parastarfeizabadi, M. & Kouzani, A.Z., 2017. Advances in closed-loop deep brain stimulation devices. Journal of neuroengineering and rehabilitation, 14(1), p.79.

Park, J. et al., 2013. Screening fluorescent voltage indicators with spontaneously spiking HEK cells. PLoS ONE, 8(12), pp.1–10.

Park, S. Il et al., 2016. Stretchable multichannel antennas in soft wireless optoelectronic implants for optogenetics. Proceedings of the National Academy of Sciences, 113(50), pp.E8169–E8177. Available at: http://www.pnas.org/lookup/doi/10.1073/pnas.1611769113.

Piech, D.K. et al., 2020. A wireless millimetre-scale implantable neural stimulator with ultrasonically powered bidirectional communication. Nature Biomedical Engineering, 4(February), pp.207–222. Available at: http://dx.doi.org/10.1038/s41551-020-0518-9.

Pinnell, R.C., Vasconcelos, A.P. De & Cassel, J.C., 2018. A Miniaturized, Programmable Deep-Brain Stimulator for Group-Housing and Water Maze Use. Frontiers in Neuroscience, 12(April), pp.1–11.

Ribeiro, C. et al., 2016. Proving the suitability of magnetoelectric stimuli for tissue engineering applications. Colloids and Surfaces B: Biointerfaces, 140, pp.430–436. Available at: http://dx.doi.org/10.1016/j.colsurfb.2015.12.055.

Roy, B., C., M.D. & A., T.P., 2007. The brain tissue response to implanted silicon microelectrode arrays is increased when the device is tethered to the skull. Journal of Biomedical Materials Research Part A, 82A(1), pp.169–178. Available at: https://doi.org/10.1002/jbm.a.31138.

Sahin, M. & Pikov, V., 2011. Wireless Microstimulators for Neural Prosthetics. Critical Reviews in Biomedical Engineering, 39(1), pp.63–77.

Schabrun, S.M. et al., 2014. Targeting chronic recurrent low back pain from the top-down and the bottom-up: a combined transcranial direct current stimulation and peripheral electrical stimulation intervention. Brain Stimul, 7(3), pp.451–459. Available at: http://www.ncbi.nlm.nih.gov/pubmed/24582372.

Seo, D. et al., 2016. Wireless Recording in the Peripheral Nervous System with Ultrasonic Neural Dust. Neuron, 91(3), pp.529–539. Available at: http://dx.doi.org/10.1016/j.neuron.2016.06.034.

Shin, G. et al., 2017. Flexible Near-Field Wireless Optoelectronics as Subdermal Implants for Broad Applications in Optogenetics. Neuron, 93(3), p.509–521.e3.

So, R.Q., Mcconnell, G.C. & Grill, W.M., 2017. Frequency-dependent, transient effects of subthalamic nucleus deep brain stimulation on methamphetamine-induced circling and neuronal activity in the hemiparkinsonian rat. Behavioural Brain Research, 320, pp.119–127. Available at: http://dx.doi.org/10.1016/j.bbr.2016.12.003.

Summerson, S.R., Aazhang, B. & Kemere, C.T., 2014. Characterizing Motor and Cognitive Effects Associated With Deep Brain Stimulation in the GPi of Hemi-Parkinsonian Rats. IEEE Transactions on Neural Systems and Rehabilitation Engineering, 22(6), pp.1218–1227.

Sun, Y. et al., 2017. Wirelessly Powered Implantable Pacemaker with On - Chip Antenna. IEEE, pp.1242–1244.

Theodore, W.H. & Fisher, R.S., 2004. Review Brain stimulation for epilepsy. The Lancet, 3(February), pp.111–118.

Ting, J.T. et al., 2014. Acute brain slice methods for adult and aging animals: application of targeted patch clampanalysis and optogenetics. Methods in molecular biology (Clifton, N.J.), 1183, pp.221–242. Available at: http://www.ncbi.nlm.nih.gov/pmc/articles/PMC4219416/.

Wan, C. & Bowen, C.R., 2017. Multiscale-structuring of polyvinylidene fluoride for energy harvesting: the impact of molecular-, micro- and macro-structure. J. Mater. Chem. A, 5(7), pp.3091–3128. Available at: http://xlink.rsc.org/?DOI=C6TA09590A.

Yu, Z. et al., 2020. An 8.2mm 3 Implantable Neurostimulator with Magnetoelectric Power and Data Transfer. In International Solid-State Circuits Conference. pp. 6–8.

Yue, K. et al., 2012. Magneto-Electric Nano-Particles for Non-Invasive Brain Stimulation. PLoS ONE, 7(9), pp.1–5.

Zhai, J. et al., 2006. Giant magnetoelectric effect in Metglas/polyvinylidene-fluoride laminates. Applied Physics Letters, 89(8), pp.8–11.

